# Histone H3 lysine methyltransferase activities control compartmentalization of human centromeres

**DOI:** 10.1101/2025.07.01.662447

**Authors:** Pragya Sidhwani, Annie Trinh, Jacob P. Schwartz, Kelsey A. Fryer, Aaron F. Straight

**Affiliations:** Department of Biochemistry, Stanford University School of Medicine; Department of Molecular and Cellular Physiology, Stanford University; Department of Genetics, Stanford University

## Abstract

Eukaryotic genomes are partitioned into discrete, functional chromatin domains, yet how adjacent domains are maintained as separate units remains unclear. Centromeres exemplify this functional separation with a kinetochore forming domain specified by the histone H3 variant CENP-A surrounded by domains of heterochromatin. Until recently, understanding this functional separation was hindered by incomplete genome assemblies and the difficulty of mapping repetitive sequences. Here, we develop approaches to map H3K9me2, H3K9me3 and CENP-A across complex centromeric repeat domains after disrupting the human H3K9 methyltransferases, SUV39H1, SUV39H2, and SETDB1, that establish heterochromatin in humans. We find that SETDB1 promotes H3K9me2 deposition, whereas SUV39H1/2 catalyze H3K9me2-to-H3K9me3 conversion at the CENP-A-rich core. Loss of all three enzymes restores H3K9me3 through a non-canonical G9a/GLP-dependent pathway while promoting CENP-A expansion into pericentromeres. Centromere adaptation maintains chromosome segregation fidelity after methyltransferase loss, revealing remarkable plasticity in centromere epigenetic maintenance. Importantly, CENP-A domain expansion is not explained by G9a/GLP-mediated heterochromatin deposition revealing a novel catalytic activity-independent function of SETDB1 in defining centromere domain boundaries. Our findings highlight unexpected plasticity in centromere organization and the multifaceted functions of chromatin regulators.

## Introduction

Heterochromatin plays a fundamental role in shaping and safeguarding the eukaryotic genome. By compacting chromatin into a transcriptionally repressive state, it helps regulate gene expression, suppress harmful recombination events, silence selfish genetic elements, and organize the genome into distinct functional domains, including centromeres and telomeres^1,2^. Broadly, heterochromatin exists in two forms: facultative and constitutive. Facultative heterochromatin is dynamic—it forms during development as cells acquire specialized identities, tailoring gene expression to support cell-type-specific functions. In contrast, constitutive heterochromatin is stably maintained across most cell types and throughout the life of the organism, acting as a silencing mechanism for repetitive elements and other potentially destabilizing genomic regions. Together, these two forms of heterochromatin coordinate genome stability and functionality.

The establishment of constitutive heterochromatin relies on a cascade of chromatin modifications, most notably the methylation of histone H3 on lysine 9 (H3K9). This mark, particularly in its trimethylated form (H3K9me3), serves as a binding platform for heterochromatin protein 1 alpha (HP1α), which helps compact chromatin into a transcriptionally silent state^1,2^. Three histone methyltransferases are central to this process: SETDB1, which can catalyze mono-, di-, and tri-methylation of H3K9, and SUV39H1/SUV39H2, which specifically promote H3K9 trimethylation but can also perform dimethylation^3^ for subsequent trimethylation by SUV39H1/2 and SETDB1. Together, these enzymes coordinate the formation of dense heterochromatin at repetitive elements, ensuring their repression and structural insulation. When this system is disrupted, repetitive elements—including transposable elements—can become reactivated, latent viruses may re-emerge, and nuclear organization is compromised. Such defects can destabilize the genome, increasing the risk of aneuploidy and contributing to tumorigenesis^2,4,5^.

Much of the constitutive heterochromatin is concentrated at centromeric regions, which are large contiguous domains of repeat sequences comprising greater than 6% of the human genome^6,7^. Centromeres can be broadly divided into two domains: core centromeres and pericentromeres. Core centromeres are primarily composed of 171 bp α-satellite sequences that are tandemly repeated to form a homogenous sequence termed the higher order repeat (HOR)^8^. They are critical for accurate chromosome segregation during mitosis and meiosis, serving as the assembly site for the kinetochore, a multiprotein complex that mediates attachment to spindle microtubules^9^. The flanking pericentromeres also contain α-satellite repeats; however, because these repeats are evolutionarily older and structurally divergent, they are called divergent HOR (dHOR) and monomeric repeats (mon)^10^. Pericentromeres play a key role in maintaining sister chromatid cohesion, ensuring proper chromosome alignment and timely separation^11–13^. In this way, both core- and peri-centromeres are important for chromosome segregation during cell division.

Apart from α-satellite sequences, both core- and peri-centromeres are interspersed with many other repeat elements, including but not limited to LINEs and SINEs^6^. Although they share similar sequence content, core- and peri-centromeres (collectively referred to as “centromeres”) are epigenetically distinct. Whereas both domains have high levels of constitutive heterochromatin^14,15^, core centromeres are unique in that they possess nucleosomes containing the histone H3 variant CENP-A that determine the site of kinetochore formation for chromosomal segregation. Within the core centromere, much of CENP-A occupies regions of low CpG methylation, called the “centromere dip region” (CDR)^6,7^. Our prior work showed that, while one in every four nucleosomes contains CENP-A within the CDR, only one in twenty CENP-A-containing nucleosomes is found in the rest of the HOR array^14^. Thus, centromeres maintain two kinds of epigenetic boundaries: one between regions of high and low CENP-A within the HOR, corresponding to a DNA methylation boundary, and another between the HOR and the divergent sequences surrounding it, where CENP-A is almost entirely absent.

The boundaries that separate the CENP-A occupying core centromere, the pericentromere, and euchromatin of the chromosome arms must be stably maintained to avoid loss of functional centromeres and pericentromeres. In the yeast *Schizosaccharomyces pombe*, strong transcription of tRNA genes that flank pericentromeric heterochromatin prevents the spread of heterochromatin to the chromosome arms, as well as into CENP-A occupying domains^16–19^. However, whether heterochromatin actively maintains and constrains CENP-A localization to the core centromere is less well understood. Prior work in *S. pombe* suggests that SUV39H-directed deposition of heterochromatin is important for CENP-A deposition^20^. However, once established, this CENP-A chromatin is able to propagate without flanking heterochromatin, suggesting that kinetochore function does not rely upon heterochromatin after the initial establishment of the CENP-A domain.

Recent studies in human cells have implicated both heterochromatin and DNA methylation in maintaining CENP-A domain boundaries. Depletion of the H3K9 methyltransferases SETDB1 or SUV39H1/2 leads to CENP-A spreading beyond the CDR during long-term culture, and loss of the facultative heterochromatin regulator PRC2, which further reduces H3K9me3 levels, exacerbates this phenotype, suggesting that multiple heterochromatin pathways converge on CENP-A boundary maintenance^21^. Independently, DNA methylation has also been shown to restrict CENP-A expansion. Whereas rapid loss of DNA methylation causes chromosome segregation errors, progressive loss, as in an immunodeficiency, centromeric heterochromatin instability and facial anomalies (ICF) model, permits adaptation to centromeric disruption^22^. Together, these findings raise several important questions: whether boundary maintenance is mediated by H3K9 methylation itself or by non-catalytic functions of the methyltransferases; how cells adapt to large-scale changes in centromeric chromatin; and if heterochromatin and DNA methylation pathways cooperate to constrain CENP-A occupancy.

Here, we use CRISPR-based gene disruption to remove SUV39H1, SUV39H2, and SETDB1 individually and in combination, revealing distinct contributions of these enzymes to centromeric chromatin organization. Using a novel analytical pipeline to map H3K9me2, H3K9me3, and CENP-A across distinct centromeric repeat domains, we find that SETDB1 promotes H3K9me2 accumulation at centromeric α-satellite repeats, whereas SUV39H1/2 are required for efficient H3K9 trimethylation. Unexpectedly, combined loss of all three enzymes triggers a compensatory restoration of H3K9me3 at core centromeres, which we show is driven by a non-canonical activity of G9a/GLP methyltransferases not normally required for centromeric H3K9me3 deposition. This ectopic H3K9me3 is accompanied by expansion of CENP-A into adjacent pericentromeric chromatin – yet despite this altered architecture, adapted triple-knockout cells restore chromosome segregation fidelity. Beyond its catalytic role, we show that catalytically inactive SETDB1 is sufficient to suppress CENP-A spreading despite persistent defects in canonical H3K9 methylation. These mutants also exhibit elevated DNA methylation at centromeric repeats, implicating a methyltransferase-independent function for SETDB1 in centromere boundary maintenance, potentially through interactions with the DNA methylation machinery. Together, these findings define a hierarchy of H3K9 methyltransferase activities at human centromeres, uncover a mechanism of adaptive epigenetic compensation, and identify an unexpected non-catalytic role for SETDB1 in maintaining CENP-A domain boundaries.

## Results

### SUV39H and SETDB1 collaboratively regulate heterochromatin at centromeric regions

Previous studies suggest that SUV39H1/2 enzymes are responsible for depositing the H3K9me3 modification at pericentromeric regions in humans and other mammals^23,24^. The recent complete sequencing of all human centromeres showed that pericentromeric regions are replete with transposable elements (TEs) which, in chromosomal arms, are repressed by a different histone methyltransferase, SETDB1^6,25,26^. To assess the different roles of SUV39H and SETDB1 histone methyltransferases at various repeat types in centromeric regions, we created a computational framework to confidently map heterochromatin associated histone modifications at various repeat elements within their diverse centromeric contexts (Materials and Methods, Fig 1A)^27^. Specifically, we performed CUT&RUN^28^ for H3K9me3 and H3K27me3 in both SUV39H1/2 double knockout (SUV39H DKO) as well as SETDB1 knockout (SETDB1 KO) cells. We aligned the reads to the T2T-CHM13v2.0 genome^29^ uniquely with no mismatches, and summed reads that aligned to specific repeat types (as annotated by RepeatMasker)^30^ within T2T defined centromeric regions^6,25,26^. So, for example, if a read aligned to an α-satellite repeat or a human-specific LINE 1 (L1Hs) repeat within the HOR or an older monomeric repeat region of any centromere, it was labeled “ALR/Alpha_hor” or “L1Hs_mon”, respectively (Fig 1A, Materials and Methods).

**Figure 1:**
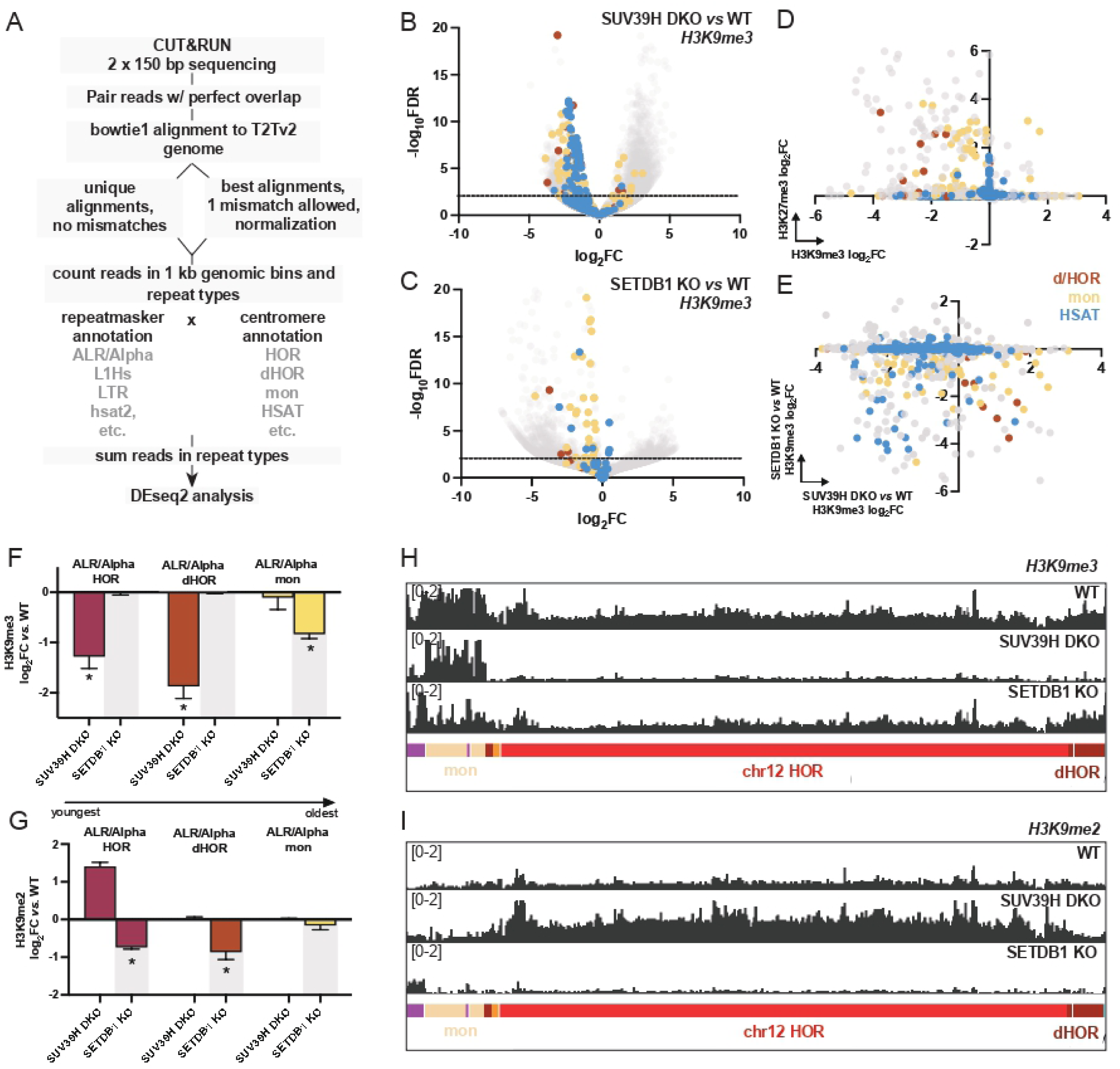
SUV39H and SETDB1 methyltransferases regulate heterochromatin deposition at centromeric regions. (A) Pipeline to process short-read CUT&RUN data from control and treated samples, for statistical comparison with DEseq2. (B-C) Volcano plots show the -log_10_ False Discovery Rate (-log_10_FDR) on the Y axis and the log_2_ fold change (log_2_FC) on the X axis for repeat elements embedded in HOR/dHOR (brown), monomeric repeat regions (yellow) and HSAT repeat regions (blue) and 50,000 other randomly sampled repetitive or genomic loci (grey) for SUV39H DKO vs. wild-type (B) and SETDB1 KO vs. wild-type (C). (D) Combined log_2_FC for SUV39H DKO or SETDB1 KO compared with their respective wild-type controls shows that H3K27me3 (Y axis) is upregulated in many of the same repeat types where H3K9me3 (X axis) is downregulated. (E) log_2_FC of H3K9me3 in SETDB1 KO vs. wild-type (Y axis) and SUV39H DKO vs. wild-type (X axis) shows that many of the repeat types that lose H3K9me3 upon SETDB1 depletion also do so upon SUV39H depletion. (F-G) A comparison of log_2_FC in H3K9me3 compared to wild-type at ALR/Alpha repeats embedded in HORs (magenta), dHORs (brown) and monomeric repeats (yellow) show that H3K9me3 is downregulated at HORs and dHORs in SUV39H DKO and at monomeric repeats in SETDB1 KO (F). On the other hand, H3K9me2 is upregulated at HORs in SUV39H DKO but downregulated in SETDB1 KO (G) (padj<0.05). (H-I) CUT&RUN tracks for H3K9me3 (H) and H3K9me2 (I) at the chr12 HOR, dHOR and monomeric regions demonstrate differences in in wild-type (top row), SUV39H1/2 KOs (middle row) and SETDB1 KOs (bottom row).

Utilizing this approach, we found that SUV39H enzymes are responsible for depositing H3K9me3 at human satellite (hsat) repeats in pericentromeres, akin to major satellite repeats in mice^31,32^ (Fig 1B). Additionally, we find that both SUV39H enzymes as well as SETDB1 deposit H3K9me3 on a multitude of repeat types in dHOR and mon repeat regions that abut the HOR regions^6^ (Fig. 1B-D). Interestingly, while H3K9me3 is broadly downregulated in the absence of these histone methyltransferases, the facultative heterochromatin mark H3K27me3 is significantly upregulated at pericentromeric regions when SUV39H enzymes or SETDB1 are absent (S1A-C). In particular, we find that 64.7% of the repeat types that have lower H3K9me3 in SUV39H DKO or SETDB1 KO compared to wild-type (log_2_FC < 0) accumulate compensatory H3K27me3 (log_2_FC > 0) (Fig. 1D). Together, these findings reveal a complex interplay between constitutive and PRC1/2-mediated facultative heterochromatin^33^ at pericentromeric repeats.

Given that both SUV39H enzymes and SETDB1 enzymes repress centromeric regions, we next wanted to investigate if they act independently or cooperatively at different core- and peri-centromeric repeat types. For the purpose of this analysis, we first focused on repeats embedded in HOR, dHOR, mon, human satellites, beta satellites, gamma satellites and other centromeric satellites that exhibit a log_2_FC of ±1 or greater compared to wild-type. We find that, for these repeat-rich regions in the centromere, 7.4% exhibit a dramatic reduction in H3K9me3 levels when SETDB1 is depleted, of which 50.8% repeats are also affected by SUV39H1/2 depletion (Fig. 1E). In contrast, SUV39H1/2 enzymes repressed 34.7% repeat types within these regions, of which only 10.8% are also highly repressed by SETDB1 (Fig. 1E). Additionally, SUV39H1/2 depletion paradoxically increases H3K9me3 at 4.4% of repeats, unlike SETDB1 depletion, which rarely causes such increases (0.9%) (Fig 1E). Together, these findings indicate that SUV39H enzymes have a wider independent role in repressing centromeric repeats.

### SETDB1 and SUV39H1/2 deposit H3K9me2 and H3K9me3 at centromeric higher order repeats

Given that older transposable elements are transcriptionally inert and therefore may be prone to a different silencing mechanism^25^, we tested whether α-satellite sequences in HORs, older dHORs, and even older monomeric repeats (mon) are repressed differently by SUV39H enzymes versus SETDB1. Our analysis revealed that H3K9me3 is significantly reduced at α-satellite repeats in HORs and dHORs in SUV39H1/2 DKO, but not in SETDB1 KO, suggesting that SUV39H enzymes deposit H3K9me3 at evolutionarily younger α-satellite repeats (Fig. 1F,H). In contrast, at older α-satellite repeats embedded in distal monomeric repeat regions, H3K9me3 levels were significantly reduced in SETDB1 KOs, but not SUV39H DKOs (Fig. 1F,H). Together, these data suggest that α-satellite repeats can be repressed differently based on their centromere context, and SUV39H1/2 enzymes are the primary methyltransferases depositing H3K9me3 at CENP-A-containing HORs.

Previous studies have shown that the CENP-A core is bounded by H3K9me2-defined heterochromatin^34,35^. To test if SUV39H1/2 or SETDB1 promote dimethylation of H3K9 at centromeric α-satellites, we performed CUT&RUN for H3K9me2 in SUV39H1/2 DKO or SETDB1 KO using an antibody that does not exhibit cross-reactivity with H3K9me3 in similar assays^36^. We find that in the absence of SUV39H enzymes, H3K9me2 accumulates at α-satellite regions in HORs, suggesting that SUV39H1/2 is responsible for the H3K9me2 to H3K9me3 transition at HORs (Fig. 1G,I). Surprisingly, H3K9me2 is significantly reduced at HOR and dHOR α-satellites in SETDB1 KOs, despite normal levels of H3K9me3 (Fig. 1G,I). These findings suggest that centromeres contain a distinct pool of SETDB1-dependent H3K9me2, consistent with previous reports of stable H3K9me2 at centromeric chromatin^34^. Supporting this idea, we identify subregions within HOR arrays that accumulate H3K9me2 in a SETDB1-dependent manner without corresponding enrichment of H3K9me3 (Fig. S1D).

While both young and old LINEs are present in dHORs and mon regions, only the evolutionarily youngest and transposition-competent LINEs, L1Hs elements, are found interspersed with HOR α-satellites^6^. Depletion of components of the HUSH complex, which targets SETDB1 to its genomic loci of repression, causes an upregulation of L1Hs transcripts^25^. We therefore wondered if SETDB1 would repress L1Hs in the context of SUV39H-regulated centromeric chromatin. To test this, we assayed H3K9me3 in L1Hs elements embedded in HORs, dHORs and monomeric repeat regions. Our data show that SUV39H enzymes deposit H3K9me3 on L1Hs elements embedded in HORs and dHORs. Therefore, repression of L1Hs mirrors that of the α-satellites they are surrounded by, suggesting that SUV39H- and SETDB1-mediated repression is driven not by the genetic identity of the underlying repeat element, but by its chromatin context (Fig. S1E).

### Depletion of SUV39H1, SUV39H2 and SETDB1 histone methyltransferases restores centromeric heterochromatin

Our findings indicate that SETDB1 deposits H3K9me1/2 whereas SUV39H methyltransferases deposit H3K9me3 at HORs. We therefore hypothesized that depleting all three methyltransferases will abolish constitutive heterochromatin at CENP-A-containing HORs. To test this, we isolated a clonal population where all three histone methyltransferases were knocked out (SETDB1/SUV39H TKO, also referred to as TKO). Specifically, we knocked out SETDB1 methyltransferase in a clonal population of cells where both SUV39H methyltransferases had been depleted. Surprisingly, we found that H3K9me3 levels were restored at HOR regions in the TKO compared to SUV39H1/2 DKO (Fig. 2A,B). In fact, DEseq2 analysis suggested that H3K9me3 was significantly higher at HORs in TKOs compared to wild-type (Fig. 2A). This kind of upregulation appeared specific to HORs, since at dHORs, H3K9me3 was still significantly lower than wild-type (Fig. 2A), albeit not as drastically as in SUV39H1/2 DKOs (Fig. 2A, Fig. 1F).

**Figure 2:**
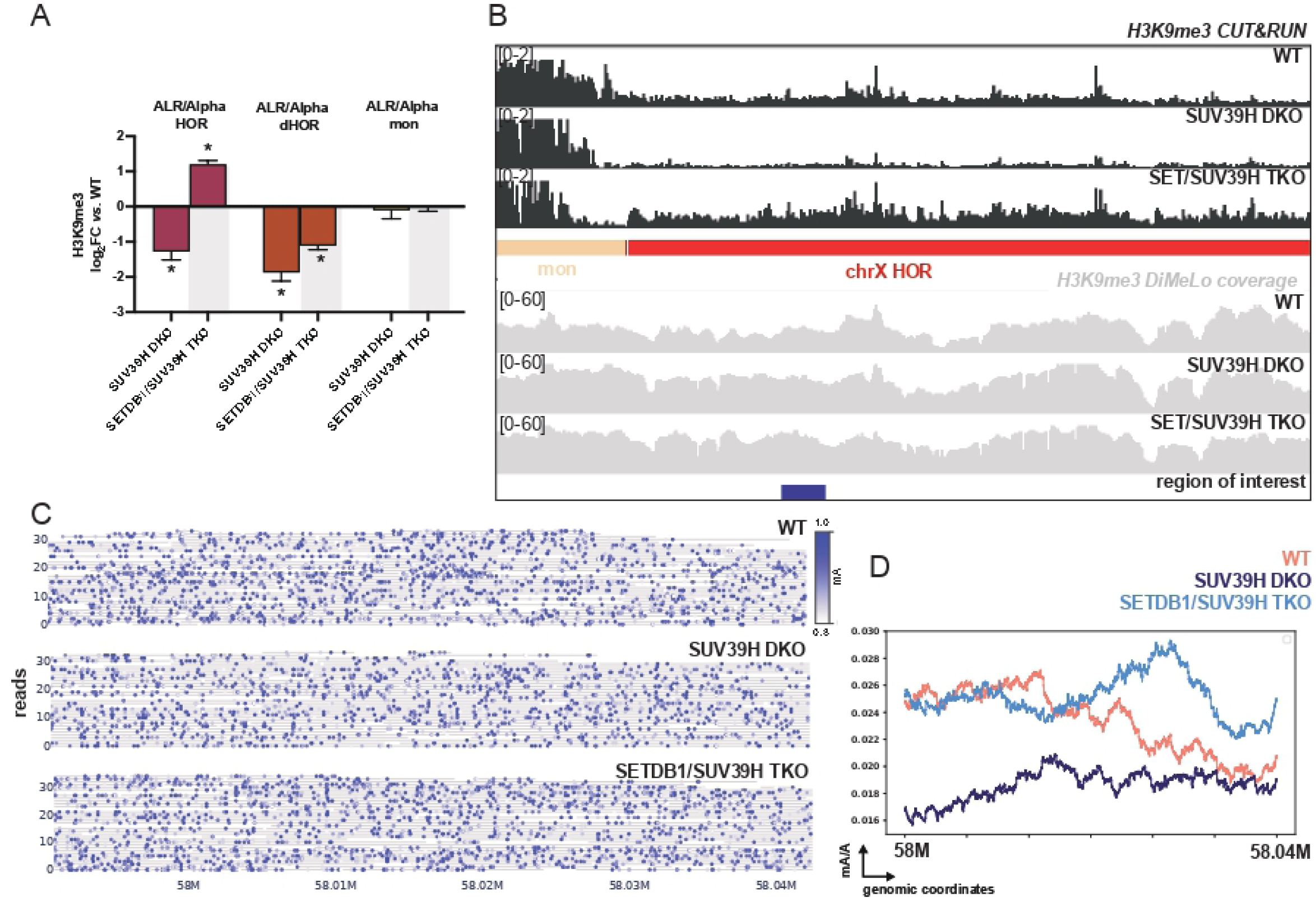
SETDB1/SUV39H TKOs have paradoxically high H3K9me3 at HORs. (A) H3K9me3 log_2_FC compared to wild-type (WT) for SUV39H KO (reproduced from Fig. 1) and SETDB1/SUV39H TKOs shows that TKOs have surprisingly high H3K9me3 at ALR/Alpha repeats embedded in HORs (magenta) but not those in dHORs (brown) or monomeric repeats (yellow) (padj<0.05). (B) CUT&RUN tracks for H3K9me3 on the chrX HOR in wild-type (1^st^ row), SUV39H DKO (2^nd^ row) and SETDB1/SUV39H TKO (3^rd^ row) show that H3K9me3 is restored in TKOs specifically at HORs. DiMeLo-seq coverage tracks for H3K9me3 for wild-type (4^th^ row), SUV39H DKO (5^th^ row) and SETDB1/SUV39H TKO (6^th^ row) show that we get sufficient sequencing depths for all conditions at regions of interest (blue, 7^th^ row). (C) Single fiber browser tracks for H3K9me3-directed methylation of adenines shows that methyl-A (mA) density (blue gradient) is higher in WT (top) and TKOs (bottom) compared to SUV39H DKO (middle) at at the region of interest highlighted in (B). (D) Line plots of mA/A for WT (salmon), SUV39H DKO (navy blue) and SETDB1/SUV39H TKO (light blue) at the region of interest (B) display that TKOs have higher H3K9me3 compared to wild-type and SUV39H DKO.

To validate these observations, we next used H3K9me3-directed DiMeLo-sequencing followed by ultra-long Oxford Nanopore sequencing to map the locations of methylated histones^14^. We focused our analysis on the HORs of three chromosomes, chr8, chr12 and chrX, to obtain higher coverage at specific centromeres. We mapped the ultra-long sequences to the CHM13-T2Tv2.0 genome^29^ and retained only high-quality primary alignments for our analysis (Materials and Methods). Although mapping K562 reads to the CHM13 reference genome resulted in reduced coverage across HOR arrays due to sequence divergence at repetitive centromeric regions, coverage within portions of the HORs remained sufficient for robust analysis, and we focused our quantification on these conserved regions (Fig. 2C,D). As with CUT&RUN, we observed a considerable reduction in H3K9me3 at CENP-A occupying HORs in SUV39H1/2 DKOs, which is surprisingly rescued in the TKOs (Fig. 2C,D). Together, our data suggest that, when all three histone methyltransferases are depleted, H3K9me3 is paradoxically restored at CENP-A containing HORs.

We next focused on H3K9me2 and H3K27me3 levels in TKOs. We find that restoration of H3K9me3 levels at HORs is accompanied by H3K9me2 rescue as well (Fig. S2A). However, H3K9me2 levels are not as high in TKOs as they are in SUV39H DKOs (Fig. S2A). Additionally, H3K27me3 is also highly upregulated at HORs as well as pericentromeric regions in TKOs (Fig. S2B). In contrast, H3K4me2, a mark of active transcription, is reduced, although not significantly (padj = 0.439) (Fig. S2C). These findings suggest that when the primary histone methyltransferases that maintain centromeric heterochromatin are perturbed, the HOR acquires SETDB1/SUV39H-independent heterochromatin, with high levels of histone modifications observed in constitutive and facultative heterochromatin.

### Compensatory centromeric H3K9 methylation by G9a/GLP in SETDB1/SUV39H triple knockouts

Given the unexpected restoration of H3K9me3 upon depletion of both SUV39H1/2 and SETDB1, we sought to identify the methyltransferase(s) responsible for H3K9 methylation in their absence. The histone methyltransferases G9a (EHMT2) and GLP (EHMT1) function as a heteromeric complex to mono- and di-methylate H3K9, primarily in euchromatin^37–39^. To test if they deposit H3K9me3 at HORs, we treated wild-type or SETDB1/SUV39H1/2 TKOs with UNC0638, a potent and specific inhibitor of G9a/GLP methyltransferase activity^40^ (Fig. 3A).

**Figure 3.**
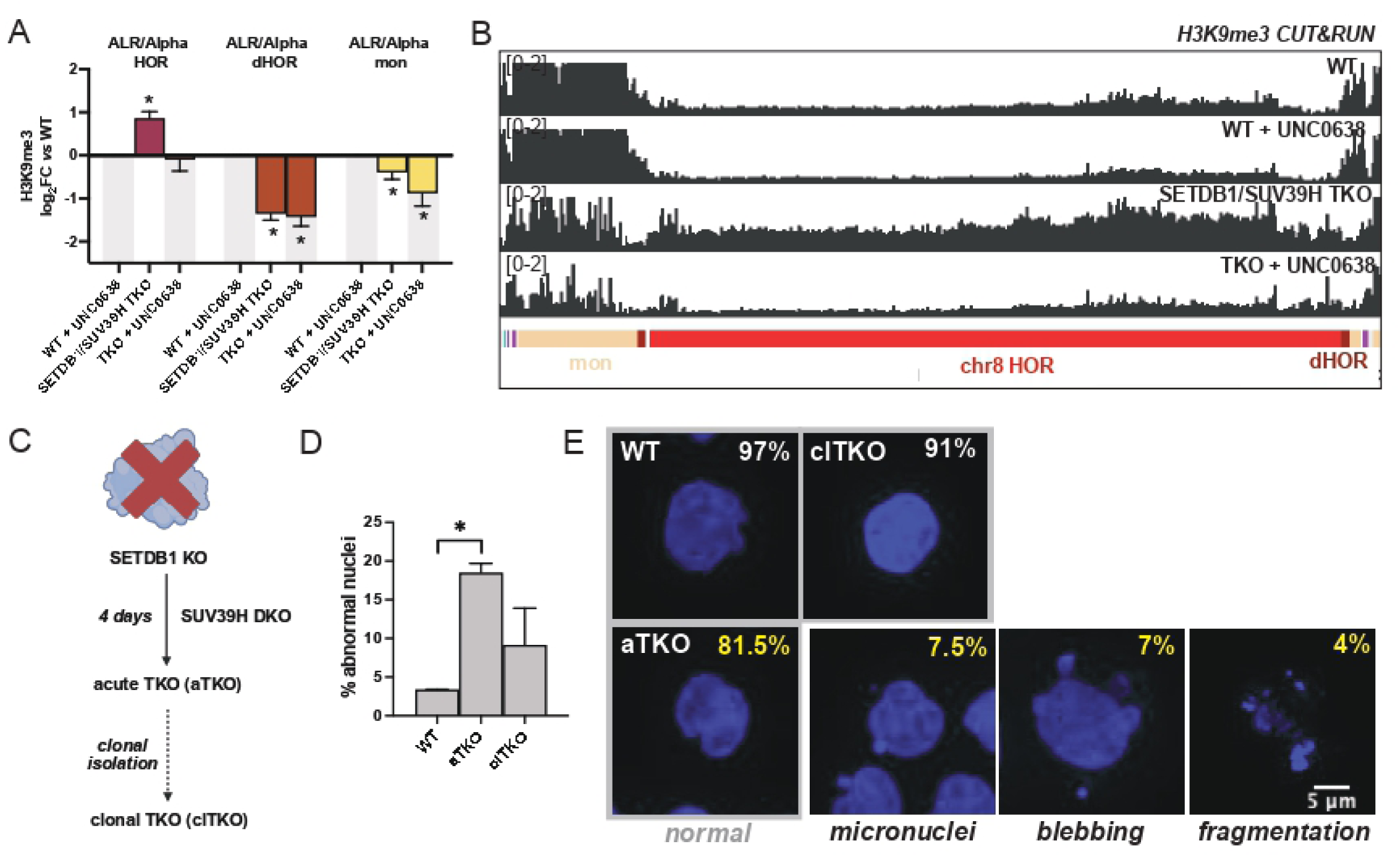
Clonal SETDB1/SUV39H TKOs exhibit G9a/GLP-mediated heterochromatin and rescued chromosome segregation. (A) H3K9me3 levels in wild-type (WT) cells remain unchanged upon UNC0638 treatment across all centromeric repeat classes. In contrast, TKOs exhibit significantly elevated H3K9me3 levels at HOR ALR/Alpha repeats (magenta; padj < 0.05), which are reduced to WT levels following UNC0638 treatment. Levels at dHORs (brown) and monomeric α-satellite repeats (yellow) remain largely unchanged.(B) Representative H3K9me3 CUT&RUN tracks at the chr8 HOR. TKOs (third row) display elevated H3K9me3 levels relative to WT cells (first row), which are reduced to WT levels following UNC0638 treatment (fourth row). UNC0638 treatment has little effect on H3K9me3 levels in WT cells (second row). (C) Experimental design for chromosome segregation analyses. Acute TKOs (aTKOs) were generated by depletion of SUV39H1/2 in SETDB1 knockout cells, whereas clonal TKOs (clTKOs) were established through long-term clonal isolation and expansion. (D) Acute depletion of SUV39H1/2 in SETDB1 knockout cells results in a significantly increased frequency of abnormal nuclei compared to mock-transfected controls (p = 0.0063, unpaired two-tailed t-test). In contrast, clTKOs do not exhibit elevated levels of nuclear abnormalities. (E) Representative Hoechst-stained nuclei from WT, aTKO, and clTKO cells. Grey boxes indicate nuclei with normal morphology, whereas examples of chromosomal abnormalities, including micronuclei, nuclear blebbing, and nuclear fragmentation, are shown.

H3K9me3 levels remained unaffected at α-satellite regions in HORs, dHORs and mon in wild-type cells treated with UNC0638, suggesting that normally, G9a/GLP methyltransferases do not deposit H3K9me3 at centromeres (Fig. 3B). In contrast, UNC0638 treatment, as well as acute depletion of G9a/GLP methyltransferases, reduced H3K9me3 levels in SETDB1/SUV39H1/2 TKOs. Notably, this effect was specific to HORs and not observed at pericentromeric regions (Fig. 3A,B, Fig. S3A,D). In contrast, acute depletion of the SETDB1 paralog SETDB2 had no effect (Fig. S3A,D). Surprisingly, acute inhibition of G9a/GLP, while reducing H3K9me3, markedly increased H3K9me2 at HORs (Fig. S3B), but H3K27me3 was largely unaffected (Fig. S3C). While the enzymatic source of H3K9me2 in UNC0638-treated TKOs remains unclear, these results are consistent with altered H3K9 methylation dynamics at centromeric repeats when SUV39H/SETDB1 H3K9me3 pathways are disrupted. Together, these data demonstrate that G9a/GLP methyltransferases deposit H3K9me3 at HORs when both SUV39H1/2 and SETDB1 are depleted, to specifically preserve centromeric heterochromatin.

### Adaptation to SUV39H1/2 and SETDB1 loss mitigates chromosome segregation defects

Next, we wondered if loss of SUV39H1/2 and SETDB1 disrupts chromosome segregation during cell division. To test this, we acutely depleted SUV39H1/2 in a SETDB1 KO (Fig. 3C). Specifically, we used two CRISPR gRNAs spanning a splice site in both SUV39H1 and SUV39H2, resulting in >98% knockout efficiency within four days, as assessed by Sanger sequencing (Fig. S4A-C). Following knockout, we assessed chromosome segregation defects and genomic instability by staining DNA with Hoescht. Whereas 97% of nuclei displayed normal morphology in mock-transfected wild-type cells, acute TKOs exhibited significantly elevated frequencies of abnormal nuclei, including micronuclei, nuclear blebs, and chromosome fragmentation, which were observed in 18.5% of nuclei examined (Fig. 3D-E).

To determine whether cells adapt to the loss of SETDB1 and SUV39H1/2 over time, we next examined nuclei in the clonal TKO population, in which G9a/GLP methyltransferases restore core centromeric heterochromatin levels. Interestingly, 91% of nuclei in clonal TKOs appeared morphologically normal, although we noted some variability (Fig. 3D-E). Together, these findings suggest that acute loss of SUV39H1/2 in SETDB1-deficient cells initially compromises chromosome segregation and nuclear structure, but that these defects are substantially reduced in established clonal TKOs, consistent with cellular adaptation to altered heterochromatin states. Notably, this adaptive response parallels recent observations in an ICF syndrome model, where cells tolerate the progressive loss of centromeric CpG methylation over extended culture^22^. Collectively, these findings suggest that centromeres possess a remarkable capacity to maintain chromosome segregation despite substantial epigenetic remodeling.

### SUV39H and SETDB1 methyltransferases constrain CENP-A localization

In humans, CENP-A chromatin is flanked by domains of pericentromeric heterochromatin, but whether this heterochromatin stabilizes or constrains CENP-A localization is unclear. To test the relationship between pericentromeric heterochromatin and CENP-A chromatin, we performed CENP-A CUT&RUN in wild-type, SUV39H1/2-depleted cells and SETDB1 KOs. Given that CENP-A is assembled primarily on highly repetitive and rapidly evolving regions^6^, we used an alternative strategy to map and assess changes in CENP-A deposition. Specifically, we allowed a single mismatch in alignments of CENP-A associated reads, to account for differences between the genomes of K562 and CHM13, and normalized repeat-derived reads to the number of genomic loci they map to (Materials & Methods, Fig. 1A). Using this computational strategy, we find that the overall levels of CENP-A are unaffected between wild-type, SUV39H1/2 DKOs and SETDB1 KOs (Fig. 4B). Of note, while our assay is able to determine quantitative changes in CENP-A, we cannot assess if CENP-A peaks move within the HOR itself. Thus, our analyses do not exclude the possibility that more subtle shifts in CENP-A localization occur within HORs, as has been suggested by recent studies^21^, but rather indicate that substantial expansion of CENP-A into adjacent repeat classes is not observed under these perturbations.

**Figure 4.**
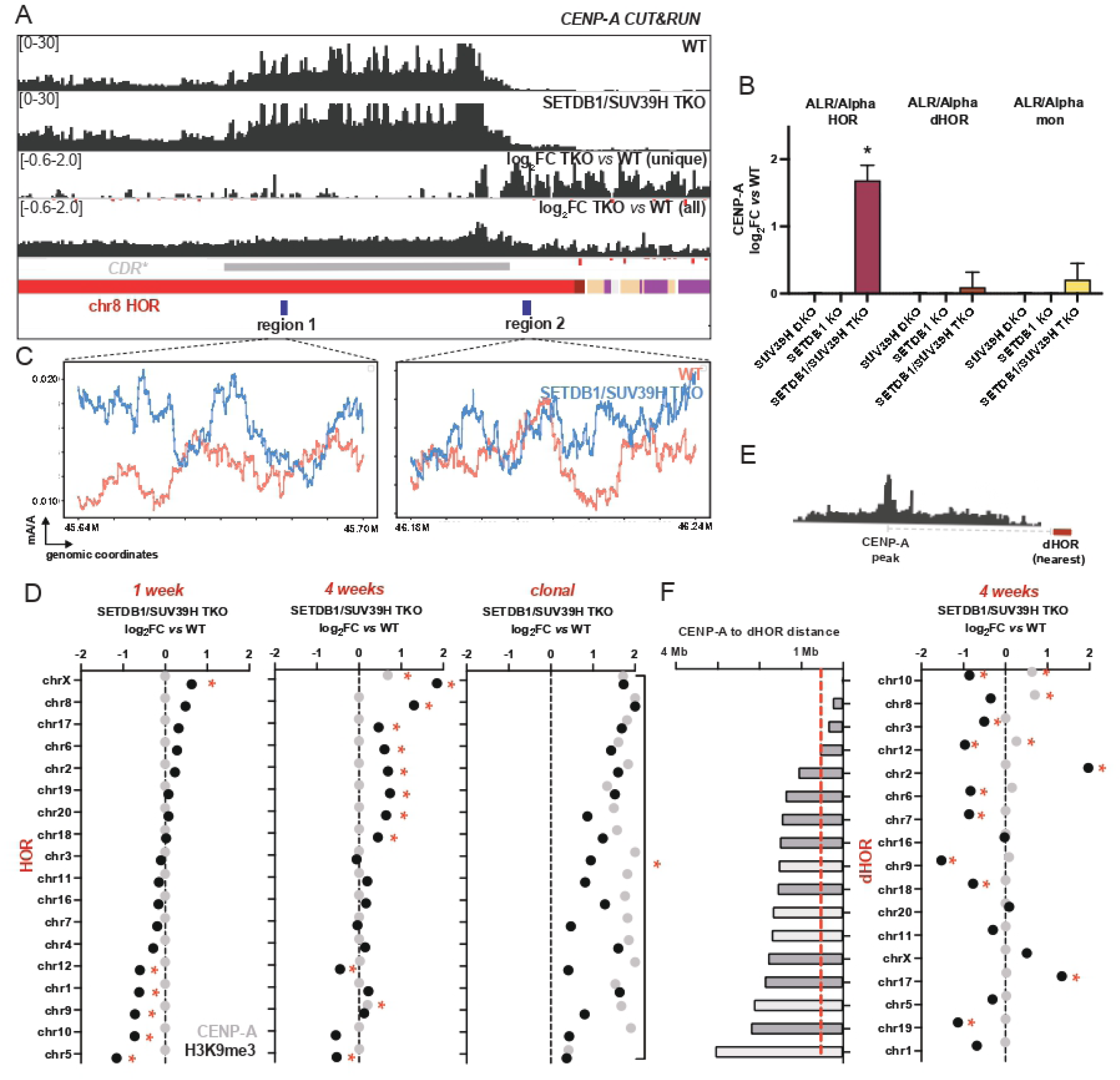
CENP-A spreads beyond its normal boundaries in SETDB1/SUV39H TKOs. (A) CENP-A CUT&RUN tracks in WT (first row) and TKO (second row) cells show increased CENP-A occupancy in TKOs. CENP-A log2FC tracks (TKO versus WT) generated using uniquely and exactly mapping reads (third row), as well as reads assigned by best alignment allowing a single mismatch (fourth row), demonstrate CENP-A spreading beyond the HOR into adjacent dHOR and monomeric α-satellite arrays. The grey bar delineates the CENP-A-defined centromere dip region (CDR*), corresponding to the region of highest CENP-A occupancy within the HOR (red). (B) CENP-A log2FC relative to matched WT controls shows little change in HOR α-satellite repeats (magenta), adjacent dHORs (brown), or monomeric α-satellite repeats (yellow) in either SUV39H DKO or SETDB1 KO cells. In contrast, SETDB1/SUV39H TKOs exhibit significantly increased CENP-A levels at HORs and modest, non-significant increases at dHORs and monomeric repeats, consistent with the spreading observed in individual centromere analyses (padj < 0.05) in (D). (C) Line plots showing the proportion of methylated adenines relative to total adenines (mA/A) within high-coverage regions (blue in A). Region 1 lies within CDR*, whereas region 2 lies outside the CDR* but within the HOR. Compared to WT cells (salmon), TKOs (light blue) exhibit CENP-A spreading toward both the p-arm (left) and q-arm (right) of the chr8 HOR. (D) Longitudinal analysis of CENP-A (grey) and H3K9me3 (black) levels at HORs across individual chromosomes one week (left) and four weeks (middle) following SUV39H depletion in SETDB1 knockout cells, as well as in clonal TKOs (right). Increases in H3K9me3 are detectable prior to substantial increases in CENP-A occupancy. In clonal TKOs, both CENP-A and H3K9me3 levels are significantly elevated relative to controls (red asterisks indicate padj < 0.05). (E) Schematic illustrating the analysis performed in (F). The distance between the WT CENP-A peak and the nearest dHOR was measured for each chromosome. (F) CENP-A (grey) expansion into dHORs is associated with proximity to the WT CENP-A peak (left bar plots) rather than with local H3K9me3 levels at individual dHORs (black). Dark grey bars indicate chromosomes for which a distinct WT CENP-A peak could be identified. For chromosomes lacking a clearly defined CENP-A peak, the distance from the center of the HOR array to the nearest dHOR was used as a proxy measurement (light grey bars).

Given the distinct nature of heterochromatin in the SETDB1/SUV39H TKOs, we tested if CENP-A spreads outside of the CDR in TKOs. Since labeling the CDR directly requires CpG methylation information, we refer to the primary CENP-A peak as the “CENP-A defined CDR”. Interestingly, we find that CENP-A levels are significantly elevated in the HOR, and slightly but not significantly higher in the dHOR and mon when all three histone methyltransferases are disrupted, suggesting that CENP-A expands outside of the CENP-A defined CDR towards pericentromeric regions when heterochromatin maintenance is disrupted (Fig. 4A, B). We further confirmed this observation by performing CENP-A directed DiMeLo-sequencing. At two regions of chr8 where we achieved sufficient coverage (>30x), one bordering the CENP-A defined CDR (region 1) and one outside of it (region 2), CENP-A levels were higher in TKOs compared to wild-type, suggesting that CENP-A is indeed spreading outwards in TKOs (Fig. 4A-C).

Importantly, CENP-A expansion was not accompanied by increased expression of CENP-A or its deposition machinery, as determined by RNA-sequencing of wild-type and triple-knockout cells (Fig. S4D).

To define the temporal relationship between heterochromatin restoration and CENP-A expansion in TKOs, we depleted SUV39H1/2 in SETDB1 KO cells and profiled H3K9me3 and CENP-A one and four weeks post-depletion. H3K9me3 levels increased above wild-type levels on a subset of chromosomes within one week, whereas CENP-A levels remained unchanged (Fig. 4D). Elevated CENP-A was first detected at the HORs of chrX and chr9 after four weeks of chronic SUV39H1/2 depletion, when half of the assayed HORs had elevated H3K9me3. Finally, in clonal TKOs, both H3K9me3 and CENP-A were significantly increased across all HORs (Fig. 4D). These findings indicate that heterochromatin restoration precedes CENP-A expansion, suggesting that the two processes are temporally separable.

Next, we asked whether CENP-A expansion into pericentromeric dHORs is driven by local loss of H3K9me3 or instead by proximity to the pre-existing CENP-A domain. To distinguish between these possibilities, we compared CENP-A occupancy and H3K9me3 levels across dHORs at varying distances from the CENP-A peak (Fig. 4E,F). In chronically depleted TKOs, CENP-A occupancy at dHORs correlated with proximity to the CENP-A peak rather than loss of H3K9me3 at individual dHORs. Specifically, CENP-A spreading extended up to ∼500 kb from the original CENP-A peak following four weeks of chronic SUV39H1/2 depletion. The sole exception was chr3, where the active HOR array is interrupted by an HSAT array^6^, placing the nearest dHOR on the opposite side of this intervening sequence. Together, these data suggest that proper heterochromatin deposition by SUV39H and SETDB1 methyltransferases is required for CENP-A boundary maintenance.

### Non-catalytic functions of SETDB1 regulate H3K9me3 accumulation at HORs in a chromosome-specific manner

How do SETDB1 and SUV39H1/2 regulate H3K9me3 and CENP-A levels at the HORs? To uncouple the methyltransferase from its methylation activity, we used adenine base editing to generate a cell line in which the catalytic activity of SETDB1 was abrogated (Materials and Methods). Specifically, we mutated cysteine 1226 within the SET domain to arginine (C1226R), since substitution of this cysteine was previously shown to abolish SETDB1 methyltransferase activity^41^. Homozygous C1226R mutants (SETDB1^m^) failed to deposit H3K9me3 at ZNF genes known to be repressed by SETDB1^41,42^ (Fig. S5A), despite maintaining wild-type levels of SETDB1 expression (Fig. S5B). These results indicate that SETDB1 catalytic activity was successfully disrupted without compromising protein stability.

We next asked whether loss of SETDB1 catalytic activity in SUV39H DKOs would recapitulate the restoration of H3K9me3 observed in TKOs. While homozygous SETDB1^m^ clones could be readily isolated in a wild-type background, we were unable to generate homozygous SETDB1^m^ clones in a SUV39H DKO background, suggesting that this combination of mutations may be deleterious. We therefore adopted an alternative strategy, performing pooled knockouts of SUV39H1 and SUV39H2 in clonal SETDB1 KO or SETDB1^m^ cell lines (Fig. 5A). We then performed H3K9me3 CUT&RUN after one and four weeks to assess the effects of acute and chronic SUV39H depletion in these backgrounds.

**Figure 5.**
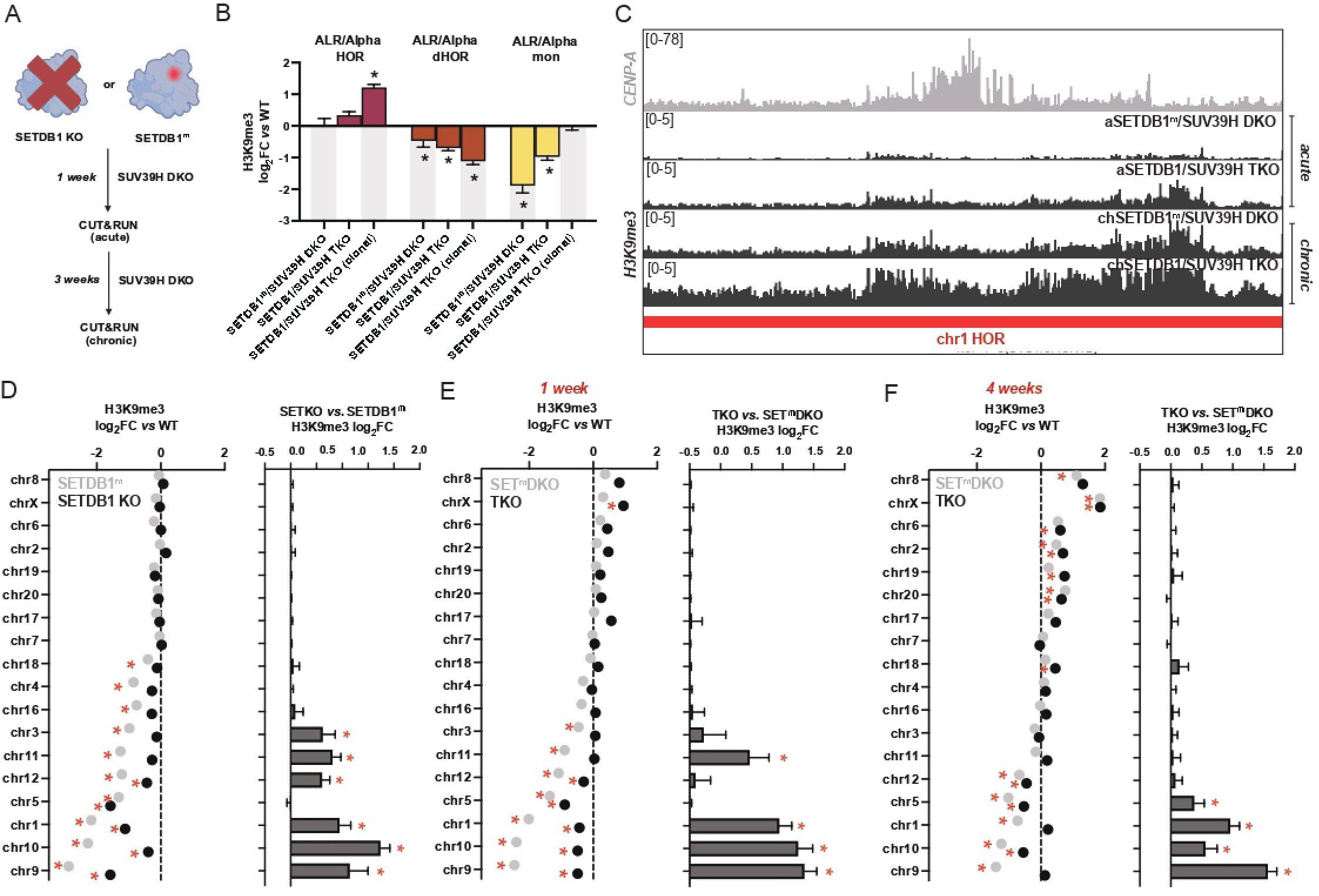
Catalytically inactive SETDB1 restrains ectopic H3K9me3 deposition at HORs. (A) Experimental schematic. SUV39H1/2 were depleted in either a SETDB1 knockout (SETDB1 KO) or catalytically inactive SETDB1 (SETDB1^m^) background. H3K9me3 CUT&RUN was performed in pooled populations after one week (acute) or four weeks (chronic) of SUV39H1/2 depletion. (B) H3K9me3 log_2_FC relative to wild-type (WT) controls in chronically depleted SETDB1^m^/SUV39H DKO pools, matched SETDB1/SUV39H TKO pools, and a clonal SETDB1/SUV39H TKO population (reproduced from Fig. 2A). ALR/Alpha repeats within HORs (magenta) exhibit significantly elevated H3K9me3 in clonal TKOs and a modest, non-significant increase in chronically depleted TKO pools relative to SETDB1^m^/SUV39H DKO pools. In contrast, all conditions show reduced H3K9me3 at dHORs (brown). (C) Representative CUT&RUN profiles at the chr1 centromere. Tracks for CENP-A (grey) and H3K9me3 (black) show reduced H3K9me3 across the HOR in SETDB1m/SUV39H DKO pools (second and fourth rows) compared to matched SETDB1/SUV39H TKO pools (third and fifth rows) following either acute (one week) or chronic (four week) SUV39H1/2 depletion. (D) Chromosome-level analysis of H3K9me3 at HORs in SETDB1^m^ (grey) and SETDB1 KO (black) cells prior to SUV39H1/2 depletion. Comparison to WT (left) and direct comparison between genotypes (right) demonstrates that SETDB1m cells exhibit significantly lower H3K9me3 than SETDB1 KO cells at a subset of chromosomes. (E,F) Chromosome-level H3K9me3 analysis one week (E) or four weeks (F) following SUV39H1/2 depletion. Reduced H3K9me3 in SETDB1^m^/SUV39H DKO cells relative to SETDB1/SUV39H TKO cells persists at a subset of chromosomes under both acute and chronic conditions. Red asterisks indicate padj < 0.05.

Across all HORs, pooled chronic TKOs exhibited a modest but not statistically significant increase in H3K9me3 relative to SETDB1^m^/SUV39H1/2 DKO cells (log_2_FC vs. WT of 0.34 vs 0.03)(Fig. 5B). However, individual HORs showed more pronounced differences. For example, H3K9me3 levels at the chr1 HOR were significantly elevated within one week of SUV39H1/2 depletion in SETDB1 KO cells compared with SETDB1^m^ cells, and this increase persisted following prolonged culture (Fig. 5C). Motivated by these chromosome-specific effects, we systematically compared H3K9me3 levels across chromosomes in SETDB1 KO and SETDB1^m^ cells before and after SUV39H1/2 depletion (Fig. 5D-F).

Surprisingly, we found that SETDB1^m^ cells exhibited significantly lower H3K9me3 levels than SETDB1 KO cells on multiple chromosomes even prior to SUV39H1/2 depletion (Fig. 5D). Importantly, many of these differences persisted following acute and chronic SUV39H1/2 depletion, indicating that loss of SETDB1 protein and loss of SETDB1 catalytic activity have distinct consequences for heterochromatin accumulation at HORs (Fig. 5E-F).

One possible explanation for this difference was that complete loss of SETDB1 protein triggered compensatory H3K9 methyltransferase activity. To test this possibility, we acutely depleted G9a/GLP or SETDB2 in SETDB1 KO cells. However, neither perturbation further reduced H3K9me3 levels (Fig. S5E). We also considered whether the catalytically inactive SETDB1 protein might remain associated with chromatin and sterically limit access of other methyltransferases. To test this idea, we performed SETDB1 ChIP-qPCR in wild-type, SETDB1^m^, and SETDB1 KO cells using primers targeting the ZNF594 locus (Materials and Methods). Contrary to this hypothesis, the catalytic mutant exhibited reduced chromatin association relative to wild-type SETDB1 at the ZNF locus (Fig. S5F). These results argue against a simple model in which persistent chromatin binding by catalytically inactive SETDB1 limits heterochromatin accumulation at HORs. However, they do not exclude more indirect mechanisms by which SETDB1 may influence heterochromatin formation. Altogether, these findings suggest that SETDB1 contributes to the regulation of HOR-associated H3K9me3 through mechanisms that are partly independent of its catalytic activity.

### SETDB1 maintains CENP-A boundaries independent of its catalytic activity

Our results demonstrate that SETDB1/SUV39H1/2 triple knockout (TKO) cells exhibit G9a/GLP-dependent heterochromatin deposition accompanied by CENP-A spreading. We next asked whether CENP-A spreading is a direct consequence of excess G9a/GLP-mediated heterochromatinization, or whether SETDB1 and SUV39H1/2 themselves contribute to the maintenance of CENP-A boundaries.

To distinguish between these possibilities, we used our pooled knockout system to deplete SUV39H1/2 in either SETDB1 KO or catalytically inactive SETDB1^m^ cells. One week after SUV39H1/2 depletion, CENP-A distribution was largely indistinguishable between the two genetic backgrounds, indicating that both begin with comparable CENP-A occupancy across centromeric domains (Fig. S6A).

We then examined the effects of chronic perturbation. Acute SETDB1/SUV39H1/2 TKOs were cultured in the presence of DMSO or the G9a/GLP inhibitor UNC0638, with the inhibitor replenished every four days. In parallel, we maintained SETDB1^m^/SUV39H1/2 double knockout (DKO) cells under matched conditions (Fig. 6A). After four weeks, we assessed CENP-A levels in TKOs, UNC0638-treated TKOs, and SETDB1^m^/SUV39H1/2 DKOs relative to wild-type cells. CENP-A spreading was readily observed in SETDB1/SUV39H1/2 TKOs and was not significantly altered by UNC0638 treatment, indicating that long-term reduction of G9a/GLP activity is not sufficient to suppress CENP-A expansion (Fig. 6B, Fig. S6B). Strikingly, however, SETDB1^m^/SUV39H1/2 DKOs exhibited substantially reduced CENP-A spreading compared to TKOs across multiple centromeric and pericentromeric repeats (Fig. 6B-C, Fig. S6C-D). In fact, DEseq2 analysis suggests that chronically depleted SETDB1^m^/SUV39H1/2 DKOs have significantly reduced CENP-A even when compared to wild-type cells on many HORs (Fig. S6D). DiMeLo-seq validation at high-coverage loci flanking the chr8 CDR confirmed increased CENP-A enrichment in TKOs relative to SETDB1^m^/SUV39H1/2 DKOs (Fig. 6D).

**Figure 6.**
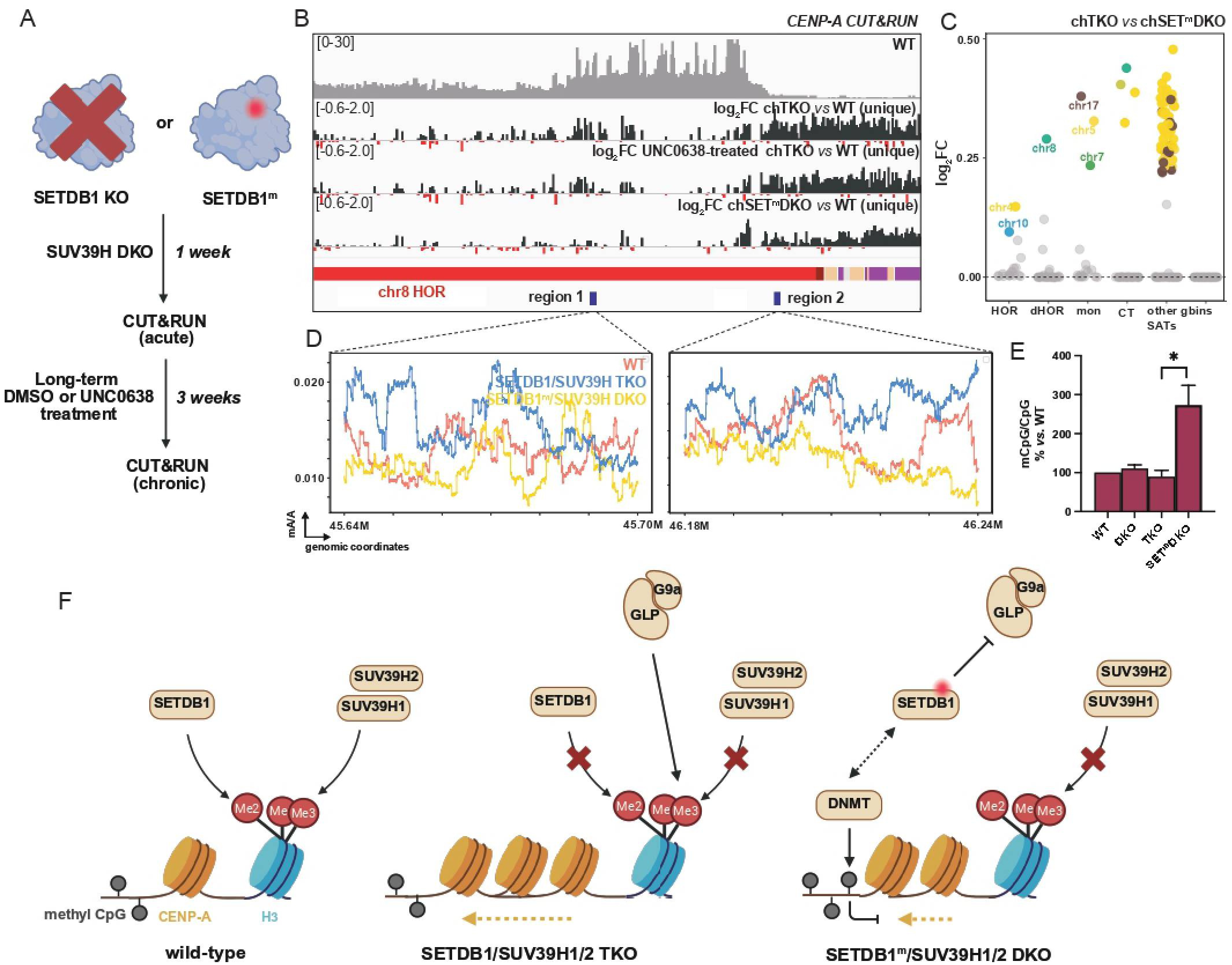
SETDB1 restricts CENP-A spreading independently of its catalytic activity. (A) Experimental design. CENP-A CUT&RUN was performed in SETDB1/SUV39H TKO pools, SETDB1^m^/SUV39H DKO pools, and UNC0638-treated SETDB1/SUV39H TKO pools to determine whether catalytic-independent functions of SETDB1 and G9a/GLP-mediated H3K9 methylation contribute to CENP-A boundary maintenance. (B) CENP-A CUT&RUN tracks in wild-type cells (gray; first row), followed by CENP-A log_2_FC relative to wild-type controls (unique alignments, no mismatches) in chronically depleted SETDB1/SUV39H TKO pools treated with DMSO (second row) or UNC0638 (third row), and SETDB1^m^/SUV39H DKO pools (fourth row). CENP-A spreads beyond the HOR in SETDB1/SUV39H TKO pools regardless of UNC0638 treatment, whereas spreading is attenuated in SETDB1^m^/SUV39H DKO pools. (C) Dot plots showing CENP-A log_2_FC in chronically depleted SETDB1/SUV39H TKO pools relative to matched SETDB1^m^/SUV39H DKO pools across repeat classes and genomic bins on individual chromosomes. Significant increases in CENP-A are observed primarily within centromeric repeat classes rather than non-centromeric genomic bins. Colored dots indicate padj < 0.05. (D) DiMeLo-seq profiles showing the fraction of CENP-A-directed methylated adenines (mA) relative to total adenines across the regions highlighted in blue in (B). These data confirm increased CENP-A occupancy in SETDB1/SUV39H TKOs (light blue) relative to wild-type (salmon) and SETDB1^m^/SUV39H DKOs (yellow). (E) CpG methylation levels measured at centromeric repeats. Bar plots show the fraction of methylated CpGs relative to wild-type cells, which were set to 100%. SETDB1^m^/SUV39H DKO cells exhibit significantly higher CpG methylation than matched SETDB1/SUV39H TKO cells (unpaired t-test, p = 0.039). (F) Model. In wild-type cells, SETDB1 promotes H3K9 dimethylation and SUV39H1/2 promote H3K9 trimethylation at HORs. In SETDB1/SUV39H TKOs, G9a/GLP mediates non-canonical H3K9 trimethylation and CENP-A expands beyond its normal boundaries. In SETDB1^m^/SUV39H DKOs, CENP-A spreading is reduced despite the presence of catalytically inactive SETDB1. These cells also exhibit elevated CpG methylation and reduced H3K9me3 at a subset of chromosomes relative to SETDB1/SUV39H TKOs, suggesting that catalytic-independent functions of SETDB1 contribute to both restriction of CENP-A spreading and suppression of G9a/GLP-mediated H3K9 methylation.

Notably, CENP-A spreading was not restricted to chromosomes where H3K9me3 accumulation was reduced in SETDB1^m^/SUV39H1/2 DKO compared to TKOs, suggesting that heterochromatin levels alone are insufficient to explain boundary disruption (Fig. 5F, Fig. 6C). We therefore considered whether other chromatin regulatory pathways contribute to the observed differences.

Given reported links between SETDB1 and DNA CpG methyltransferases^43^, and prior evidence that CpG methylation can restrict CENP-A deposition^22,43^, we profiled CpG methylation across HORs. Interestingly, SETDB1^m^/SUV39H1/2 DKOs exhibited significantly higher levels of CpG methylation across HORs compared to both wild-type and TKOs, consistent with reduced CENP-A levels (Fig. 6E). In contrast, neither SUV39H DKOs nor TKOs showed a substantial reduction in CpG methylation relative to wild-type cells (Fig. 6E).

Together, these findings indicate that SETDB1 contributes to the maintenance of CENP-A boundaries through a mechanism that is independent of its catalytic activity. The reduced CENP-A spreading observed in SETDB1^m^/SUV39H1/2 DKOs suggests that non-catalytic functions of SETDB1 are sufficient to constrain CENP-A expansion at centromeric and pericentromeric repeats. Notably, inhibition of G9a/GLP with UNC0638 does not suppress CENP-A spreading in SETDB1/SUV39H1/2 TKOs, indicating that G9a-dependent heterochromatinization does not drive CENP-A expansion. In contrast, the SETDB1^m^/SUV39H1/2 DKO state is associated with increased CpG methylation at HORs, consistent with DNA methylation influencing CENP-A boundary stability (Fig. 6F).

## Discussion

Centromeres are classically described as epigenetically bipartite, with open chromatin domains that house CENP-A, surrounded by heterochromatic pericentromeric regions that help support kinetochore function^15^. Leveraging the T2T genome assembly as well as novel approaches to map repeat-derived reads, we provide a new view of these megabase-scale regions, revealing distinct domains governed by different modes of epigenetic regulation. We uncover new roles for SETDB1 at human centromeres: it deposits H3K9me2 at younger, CENP-A-containing higher-order repeats (HORs), while also contributing H3K9me3 at older monomeric repeats. SUV39H1 and SUV39H2 build upon H3K9me2 by adding H3K9me3 to both the HORs and adjacent α-satellite regions. This two-step process likely provides finer control over heterochromatin formation at HOR regions, and may also facilitate functionally distinct roles for H3K9me2 and H3K9me3, as has been proposed in *S. pombe*^44^.

In the absence of both SETDB1 and SUV39H enzymes, G9a/GLP methyltransferases ectopically deposit H3K9me3 across CENP-A-containing HORs, indicating that these enzymes can compensate for the loss of canonical H3K9 methyltransferases at centromeres (Fig. 6F). Interestingly, this G9a/GLP-mediated heterochromatinization appears specific to HORs.

Whereas pericentromeric α-satellites exhibit reduced H3K9me3 levels in TKOs and show little response to the G9a/GLP inhibitor UNC0638, α-satellites within HORs display elevated H3K9me3 and are markedly sensitive to UNC0638 treatment. What makes HORs more susceptible to G9a/GLP-mediated heterochromatinization than dHORs, despite similar sequence content? One possibility is that distinct chromatin environments differentially recruit these complexes: H3K27me3 is substantially enriched over dHORs and monomeric repeats relative to HORs in TKOs, potentially reflecting preferential recruitment of PRC2 to pericentromeric repeats and G9a/GLP to HORs. Notably, whereas acute depletion of SUV39H1/2 in SETDB1 knockout cells results in chromosome segregation defects, clonal TKO cells adapt and regain near-normal segregation fidelity. The selective restoration of H3K9me3 at active HORs therefore raises the intriguing possibility that heterochromatin associated with the CENP-A domain plays a particularly important role in maintaining chromosome segregation.

Such a model would suggest that compensatory G9a/GLP activity preferentially safeguards the chromatin environment of the functional centromere when canonical H3K9 methyltransferases are disrupted, thereby promoting chromosome stability.

Promiscuous deposition of H3K9me3 across HORs is accompanied by CENP-A spreading in SETDB1/SUV39H TKO cells, suggesting that proper heterochromatin establishment by these enzymes is required to maintain precise CENP-A localization. Our findings provide nuanced insights into how CENP-A boundaries erode following disruption of centromeric heterochromatin. Rather than redistributing indiscriminately to regions with the greatest local loss of H3K9me3, CENP-A preferentially expands into dHORs nearest to the pre-existing CENP-A domain. These observations support a model in which CENP-A spreading occurs progressively outward from the original centromeric domain, suggesting that proximity to existing CENP-A chromatin is a major determinant of ectopic CENP-A incorporation.

Our data further indicate that the relationship between H3K9me3 and CENP-A localization is more complex than a simple antagonistic boundary model^21^. Although alterations in heterochromatin precede CENP-A redistribution during chronic depletion, stable TKO cells paradoxically accumulate elevated levels of both H3K9me3 and CENP-A at HORs. Additionally, inhibiting G9a/GLP-dependent H3K9me3 in SETDB1/SUV39H1/2 TKOs is not sufficient to curtail CENP-A spreading. Thus, H3K9me3 abundance alone appears insufficient to dictate CENP-A positioning. Instead, our observation that catalytically inactive SETDB1 retains the ability to constrain CENP-A expansion suggests that SETDB1 contributes to boundary maintenance through functions independent of its methyltransferase activity. What might these non-catalytic functions entail? SETDB1 and its paralog SETDB2 are unique amongst SET-domain methyltransferases in that they contain putative CpG binding domains. Given that HORs have high CpG methylation, a provocative idea is that SETDB1 can recognize DNA methylation, and potentially also mediate crosstalk with DNA methyltransferases, to maintain centromeric chromatin structure^45^. Consistent with this, we find that SETDB1^m^/SUV39H1/2 DKO cells exhibit increased CpG methylation at HORs together with reduced CENP-A spreading relative to TKOs, suggesting that DNA methylation-associated functions of SETDB1 may contribute to the maintenance of centromere identity. Indeed, SETDB1 directly interacts with DNMT3A and DNMT3B^43^, the DNA methyltransferases methylating centromeric regions^46–48^, suggesting regulatory feedback between DNA methylation and H3K9 methylation. While the molecular basis of this coordination remains unknown, it is interesting to note that MPP8, a protein that recruits SETDB1 for H3K9 methylation^49^, also mediates the interaction between DNMT3A and G9a/GLP methyltransferases^50^. Perhaps MPP8, SETDB1 and DNMT3 enzymes together govern DNA methylation and H3K9 methylation at HORs in a protein complex that, when SETDB1 is perturbed, is poised to recruit G9a/GLP methyltransferases.

Overall, our work underscores the modular and layered nature of epigenetic regulation at the centromere, where multiple methyltransferases act sequentially or redundantly, and where the balance between chromatin repression and CENP-A localization is tightly maintained. The differential sensitivity of HORs versus dHORs to heterochromatinization highlights an unappreciated specificity in how these enzymes interact with distinct α-satellite domains, raising fundamental questions about how sequence features, DNA methylation, and chromatin context guide the recruitment or exclusion of chromatin regulators. Finally, the interplay between DNA methylation and histone modifications at centromeres emerges as a compelling area for future exploration. Elucidating the precise molecular circuitry that maintains this balance will be critical for understanding how the epigenome defends the structural integrity of human centromeres.

## Limitations of the study

Our analyses rely on aligning sequencing reads from K562 cells to the CHM13 reference genome. Although CHM13 provides the first complete assembly of all human centromeres and enables repeat-resolved analyses that were previously impossible, centromeric α-satellite arrays are polymorphic among individuals^51^. Consequently, differences in HOR organization, repeat copy number, sequence composition, and the precise positions of CENP-A domains between K562 and CHM13 may influence the assignment and quantification of centromeric reads. To mitigate these effects, we employed mismatch-tolerant alignments and normalized repeat-derived reads by their genomic copy number to ensure that our interpretations were robust to sequence differences. Future studies incorporating haplotype-resolved assemblies of K562, particularly across the divergent CDR regions, will help to extend our findings. In addition, our conclusions regarding compensatory G9a/GLP activity are based on pharmacological inhibition and genetic perturbation rather than direct mapping of G9a/GLP occupancy. Because we were unable to obtain robust G9a/GLP CUT&RUN datasets, we cannot distinguish whether G9a/GLP are directly recruited to HORs or act through indirect mechanisms to restore H3K9 methylation following SETDB1/SUV39H disruption.

## Materials and Methods

### Cell culture

K562 cells were cultured in RPMI 1640 (Gibco 31800022) supplemented with 10% Fetal Bovine Serum (GenClone 25550) and 100U/mL Penicillin/Streptomycin (Gibco 15140163). Cells were counted with a hemocytometer every 2 days and split when they reached a density of 1 million cells/mL. The K562 cells used in this study contain a doxycycline-inducible GFP reporter and constitutively active rTetR that were introduced for independent experiments. All knockouts and cell modifications were made in this background.

### Constructing knockout cell lines

We used the CRISPOR tool (https://crispor.gi.ucsc.edu/)^52^ to design Cas9 NGG guides to target sequences that were 100-300 bp apart at the first or second intron/exon junction of SUV39H1, SUV39H2 or SETDB1. Prior work has shown that when cells are transfected with a pair of guides targeting regions <300 bp, it leads to uniform pooled knockouts with deletions of the entire region between the targets, which we could assess using PCR^53^. Two guides per protein were ordered from Synthego as single guide RNAs (sgRNA) containing 2’-O-Methyl at 3 first and last bases as well 3’ phosphorothioate bonds between first 3 and last 2 bases, which enhance gRNA stability. To construct pooled knockouts, we transfected K562 cells with two ribonucleoprotein complexes (RNP), each containing one sgRNA complexed with high fidelity Cas9 (Integrated DNA Technologies 1081060). Specifically, we added 61 pmol Cas9 to 110 pmol sgRNA, mixed by pipetting and prepared the RNP complex at room temperature for 20-30 minutes. To prepare the cells, we counted and spun 600,000 cells at 200 x g for 5 minutes, washed once with 1x Phosphate buffer saline (PBS) and aspirated the PBS. For transfection, we used the Lonza nucleofector Kit V and prepared the transfection reagent as per the manufacturer’s protocol, with 86.1 µL transfection reagent mixed with 18.9 µL supplement. We added 100 µL of this mix to the RNP complex, mixed by pipetting up and down without introducing bubbles and transferred the mix to the second RNP complex for the protein followed by mixing again. The ∼104 µL of the transfection reagent and RNP complexes was then added to cells, pipetted without bubbles and transferred to cuvettes. The cells were then nucleofected with the program T-016 in a Amaxa Nucleofector IIb device. After transfection, cells were carefully transferred to 6-well plates containing 2 mL of RPMI 1640. After 2 days, cells were transferred into flasks containing 5 mL media.

### Constructing the SETDB1 methyltransferase mutant

To make an endogenous single bp mutation in SETDB1, we used the Cas9-based A-base editing system^54^. Briefly, we designed a guide that targets the A base in the 3rd position of the reverse strand as per prior guidelines to edit it to a G. Making this A>G mutation changes the codon in the forward strand from a TGC to a CGC, which would change the cysteine in position 1226 of SETDB1 to an arginine. This guide would also likely mutate the A in the 4th position; however, that leads to a silent mutation that does not affect the amino acid encoded by that codon. Using golden gate assembly, we then cloned the guide sequence into a plasmid encoding Cas9 and the A-base editor (https://www.addgene.org/179097/). We nucleofected 1 µg of this plasmid into K562 cells then sorted single cell clones using the SONY FACS SH800S sorter and allowed them to propagate for 3 weeks. To identify cell lines with successful mutations, we PCR’d the site of mutation and performed restriction digestion with HpyCH4V (NEB Cat#R0620S), which would only digest the unedited and not the edited allele.

### Validation of knockout cell lines using PCR and Sanger sequencing

Since we used two sgRNAs per protein, we could qualitatively assess knockout efficiency using PCR. Specifically, we designed primers that bind outside the two PAM sequences and assayed for the expected reduction in PCR band size if the fragment between the two target sites was deleted. Following this, we purified the PCR reaction using the Qiagen PCR purification kit (Qiagen 28104) or using a 0.9x SPRIselect bead cleanup (BeckmanCoulter B23317) for Sanger sequencing. We then used Synthego Inference of CRISPR Edits (ICE) analysis (Synthego Performance Analysis, ICE Analysis. 2019. v3.0. Synthego) to determine the efficiency of pooled knockout^55^ (Fig. S4B-C). For SETDB1 KO and SETDB1/SUV39H TKOs, we sorted cells into single clones using the SONY SH800s FACS sorter, followed by another round of validation by PCR and Sanger sequencing on the clonal populations.

### Validation of knockout cells lines using Western Blotting

For western blot analysis of SUV39H1, we followed a previously published protocol from our lab^56^. Briefly, 10-20 million cells were washed once with 1x PBS, resuspended in 5 mL hypotonic buffer, dounced 10x with a B pestle and pelleted at 5000xg for 10 min at 4°C. Cells were then sonicated and concentration measured using a Bradford assay (Bio-Rad 5000006). 100 µg of total protein was loaded onto 17.5% SDS-PAGE gels for analysis (Fig. S4A) with 1:500 mouse anti-SUV39H1 antibody. For western blot analysis of SETDB1, G9a, GLP and SETDB2, 1 million cells were spun at 200xg for 5 minutes, washed once with 1x PBS and resuspended in 100 uL RIPA buffer containing PMSF and LPC. 15 µg of total protein was electrophoresed in 7.5% SDS-PAGE gels (Fig. S3D, Fig. S5B) and western blotted with 1:2500 dilution of antibody in Tris-buffered saline with 4% milk.

### UNC0638 treatment

UNC0638 was purchased in powder form (SelleckChem S8071) and resuspended in DMSO to a concentration of 10 mM. To test conditions, 300,000 total cells were spun at 200xg for 5 minutes and resuspended in 2 mL media containing DMSO, 250 nM UNC0638 or 500 nM UNC0638. After 4 days, cells were spun at 200xg for 5 minutes, washed with 1x PBS, and resuspended 1 mL DiMeLo Digitonin-wash buffer^14^ and exposed to an antibody against H3K9me2 at 1:100 for 2 hours. Cells were washed again and Alexa Fluor 647 antibody was added at 1:1000 for 20 minutes followed by 1:1000 Hoechst 33342 for 10 minutes. Control and treated cells were scanned using the SONY FACS SH800S. Treatment with 250 nM UNC0638 led to a decrease in 647 fluorescence but not Hoechst fluorescence, indicating that UNC0638 reduced H3K9me2 signal overall. Therefore, we used 250 nM UNC0638 as described. For long-term UNC0638 treatment for CENP-A analysis, cells were diluted in DMSO or UNC0638 containing media to 150,000 cells per mL for a total of 2 mL every four days for one month.

### CUT&RUN library preparation

CUT&RUN was performed with validated antibodies as has been previously described^28^. Specifically, 1 million cells were counted, pelleted at 200xg for 5 minutes and washed with CUT&RUN 1x wash buffer. After resuspension in the wash buffer, the cells were counted again to ensure a total of 500,000 cells per condition. Meanwhile, concanavalinA (conA) beads were activated in the binding buffer as per manufacturer’s protocol (Epicypher 211411). 10 µL of conA beads were added per condition in a 15 mL falcon tube, and the cells were shaken on a benchtop nutator for 10 minutes at room temperature. After binding the cells to the conA beads, cells were aliquoted into 500 µL LoBind tubes (Eppendorf EPPR4509), activation buffer removed, followed by resuspension in 100 µL Dig-wash buffer containing the appropriate dilution of the antibody (1:100 for anti-CENP-A, anti-H3K9me2 and anti-H3K27me3, 1:50 for anti-H3K9me3 and 1:1000 for anti-IgG). Tubes were incubated overnight at 4°C on a vortexer shaking at low speed.

The next day, cells were washed twice on the magnetic stand with 300 µL Dig-wash buffer. Cells were then incubated with 150 µL protein A fusion to micrococcal nuclease (pA-MNase) at 700 ng/µL for 1 hour at 4° C, also on a vortexer shaking at low speed. Cells were then washed twice again, followed by addition of cold Dig-wash buffer and of 2 μL 0.1 mM CaCl_2_ while shaking on a vortexer, to activate the MNase. Finally, MNase digestion was stopped by adding 100 uL of 2x STOP buffer containing 340 mM NaCl, 10 mM EDTA, and 10 mM EGTA. DNA was then extracted from cells using either ethanol precipitation or SPRIselect bead cleanup (BeckmanCoulter B23317). DNA concentrations were measured using Qubit high sensitivity reagents (Invitrogen Q32851).

For library preparation for Illumina sequencing, we used the NEBUltra II DNA library preparation kit for Illumina (NEB E7645L) following the manufacturer’s protocol. However, we edited the PCR as per the CUT&RUN protocol to do 30s of initial denaturation at 98°C, followed by 13 cycles of 10s at 98°C and 10s at 65°C, followed by a final extension of 5 minutes at 65°C and a 4°C hold. To validate size distribution of the libraries, we used high sensitivity Agilent Tapestation 4200 and if needed, conducted a final cleanup with 0.9x SPRISelect beads (BeckmanCoulter B23317). Libraries were then sequenced using 2 x 150 bp paired-end sequencing on an Illumina Novaseq X Plus platform to generate ∼50 million reads per sample.

### CUT&RUN data processing

The pipeline used for the analysis is available on GitHub (https://github.com/straightlab/cut-run_repeat_analysis/). Reads were processed by first trimming sequencing adaptors using Trimmomatic as previously described^57,58^. Trimmed reads were then paired using bbmerge with pfilter=1 to ban merges with mismatches in the overlap region^59^. The merged reads were aligned to the CHM13-T2Tv2.0 genome (https://github.com/marbl/CHM13) using bowtie1 with the tags --best --strata -X 1000 --sam-nh. We used a patched version of bowtie1 developed by Georgi Marinov that adds NH tags to the alignments. For unique alignments with no mismatches, the additional tags -v 0 -m 1 were used (Note: NH tags are not required to process these “uniqueBAMs”, so the original version of bowtie1 can be used). To report all best alignments with a single mismatch allowed, the tags -v 1 -a were used. Following this, we used a custom python script^27^ to count reads that align to various repeat types in various centromeric contexts. This script takes uniqueBAMs as input, i.e. bam files that were obtained using the -m 1 tag, or it can take bam files that have all best alignments reported, in which case it normalizes the mappings to the number of loci they map to, as per the “NH tag”. So, if one read maps equally well to 10,000 loci, each locus is assigned a weighted score of 0.0001. In addition to an indexed bam file with alignments and NH tags, this custom python script also requires a bed file to count the number of reads mapping to the regions defined in the bed file. Two types of bed files were used for this:

1. We used bedtools^60^ to intersect repeatmasker annotations^30^ with centromeric annotations (cenSatv2.0)^6^. To generate this file, we used the options -wa -wb to first generate a temporary file that contains entries from both input files. We then intersected this output to the centromeric annotation bed file to restrict the output to repeat types that are within centromeric regions. Then, we used awk commands to append the centromeric annotation from column 13 to the repeatmasker annotation in column 4 and subsequently kept only the first 7 columns. In this way, column 4 of the new bed file contains information about both the repeatmasker and the centromeric annotations for the genomic coordinates in columns 1-3. For chromosome-by-chromosome analysis, we also appended the chromosome number from column 1 to column 4. Using bedtools complement, we constructed an additional file that had repeatmasker annotations outside of the centromeric region, which we then appended to the file containing repeat information for centromeric regions. This merged file is the first type of file, used to sum read counts in repeats.
2. For DEseq2, we also wanted to input read counts for the rest of the genome not covered by any repeats. To generate a bed file for this, we used bedtools complement to remove repeat regions encoded in repeatmasker annotations from the CHM13-T2Tv2.0 genome. Following this, we used bedtools makewindows with -w 1000 to make 1000 bp windows of the genome.

Once read counts were generated for all genomic regions, a custom Python script was used to sum reads overlapping RepeatMasker centromeric annotations (column 4 of the annotation file). These repeat-level counts were then appended to the 1 kb genomic window counts. Count tables from two control and two treatment biological replicates were merged and analyzed using DESeq2 to identify differentially enriched regions. Statistical significance was determined using the Benjamini-Hochberg procedure as implemented in DESeq2, with an adjusted p-value threshold of padj < 0.05. log2 fold changes were shrunk using the ashr method^61,62^. For normalized counts of bams aligned with the -a option, the counts are not integerized, which is a requirement for DEseq2. Thus, we also integerized those counts using awk prior to DEseq2 analysis.

To generate Integrated Genome Viewer (IGV) plots of pileups, we used bam files from the -v1 -a bowtie command and processed them using a custom python script with the options -excludeReadsMappingToOtherChromosomes -RPM to generate bedgraphs of normalized counts, which were converted to bigwigs for visualization using deepTools bamCoverage^63^.

### Purification of nanobody-Hia5 fusion

The anti-rabbit nanobody-Hia5 fusion, with 6-His and MBP tags, was cloned (ATUM) into the pD861-SR vector containing Kanamycin resistance and a Rhamnose-inducible protomer. The plasmid was transformed into NEB c3010 chemically competent cells (New England Biolabs) and grown in 3L of Luria Broth at 37°C. Upon reaching OD600 of 0.6, cells were induced with 2.5mM rhamnose and grown overnight at 18°C. The pellet was thawed on ice and resuspended in 40mL of 50mM HEPES pH7.7, 300mM NaCl, 10% glycerol, 20mM imidazole, 0.5% TritonX-100, 5mM β-mercaptoethanol (β-me), with one cOmplete EDTA-free protease inhibitor tablet (Roche). Cells were lysed by tip sonication and then centrifuged at 90000xg 35k rpm for 45 minutes at 4°C. 2mL of Nickel-NTA resin (Qiagen,) pre equilibrated in binding buffer, was added and incubated at 4°C for 1 hour, after which the resin was washed thoroughly in 50mM HEPES pH7.7, 300mM NaCl, 10% glycerol, 20mM imidazole, 5mM β-me. The sample was eluted from the column in 8mL of 50mM HEPES pH7.7, 300mM NaCl, 10% glycerol, 300mM imidazole, 5mM β-me. 1mL of amylose resin (New England Biolabs), pre equilibrated in binding buffer, was added and the sample was incubated at 4°C for 90 minutes, washed in 50mM HEPES 7.7, 150mM NaCl, 10% glycerol, 1mM DTT, and eluted in 2mL of the same buffer supplemented with 20mM maltose. The eluate was supplemented with glycerol to 20% final concentration and 50uL aliquots were flash-frozen in liquid nitrogen and stored at −80C. For a detailed protocol, see: dx.doi.org/10.17504/protocols.io.6qpvrqbyzlmk/v1.

### DiMeLo-sequencing

DiMeLo-seq experiments were performed as previously described with some modifications^14^. We collected six million cells per condition, permeabilized them with 0.02% digitonin, and treated them with 1:50 rabbit (Rb) anti-H3K9me3 (Active Motif 39162) or 1:100 Rb anti-CENP-A primary antibody^64^. After washing, anti-Rb Nb-Hia5 was applied for 1hr at 250 nM. After washing off excess methyltransferase, samples were transferred to activation buffer with *S*-Adenosyl methionine and treated for 2hrs at 37°C to activate the methyltransferase activity of Hia5.

Genomic DNA was then extracted using the Monarch HMW DNA Extraction Kit for Tissue (New England Biolabs; kit code T3060). We followed the manufacturer’s guidelines with some changes, as recommended by ONT. For a detailed extraction protocol, see: dx.doi.org/10.17504/protocols.io.kqdg31eeql25/v1. Following DNA extraction, samples were prepared for sequencing following the manufacturer’s guidelines for ultra-long DNA sequencing using the SQK-ULK114 kit. After library preparation, each sample was loaded onto a promethion flowcell (FLO-PRO114M) with R10 chemistry. We used adaptive sampling to enrich for centromeric regions of chr8, 12 and X by depleting the rest of the genome. After 24 hours of sequencing, samples were extracted from the flowcells and reloaded after flowcell washing and re-priming. Total sequencing depth for each sample ranged from 10-15 gigabases.

### DiMeLo-seq data processing

Base- and modification-calling were performed on Dorado version 0.7.3 using the high accuracy model (hac) with 5mC_5hmC, and 6mA calling. Reads were aligned to the CHM13-T2Tv2.0 genome using Dorado aligner, and then quality filtered using samtools^65^ with the options -q 10 and -F 2308 to retain only high quality primary alignments. These high-quality primary read alignments are shown as coverage tracks in Fig. 2. For data visualization, we used the dimelo v2 package developed by Aaron Streets’ laboratory (https://github.com/streetslab/dimelo).

Specifically, we used the plot_enrichment_profile function to generate the line plots in Fig. 2 and Fig.6, with methylation thresholds of 0.7. We also used the plot_read_browser function to generate the single molecule plots in Fig. 2. Finally, we used the plot_enrichment function to generate aggregate CpG methylation profiles across all active HORs. While enrichment was targeted to the HORs of chromosomes 8, 12, and X, all reads aligning to active HORs were included in the analysis.

### ChIP-qPCR

ChIP was performed as described previously with some modifications^66^. Briefly, 20 million cells were washed with 1x PBS and fixed with 2 mM Disuccinimidyl Glutarate (Thermo Scientific 20593) for 45 minutes, followed by 1% formaldehyde for 20 minutes. Following nuclear extraction with buffer 1, cells were dounced 20 times with pestle A to ensure proper nuclear extraction for sonication. Cells were sonicated for 20 minutes in buffer 3 (10 mM Tris pH 8.0, 100 mM NaCl, 1 mM EDTA, 0.5 mM EGTA, 0.1% sodium deoxycholate, 0.5% N-lauroylsarcosine) on a Covaris E220 instrument with the settings Peak power = 140, Duty Factor = 10, Cycles per burst = 200. Finally, cellular debris was pelleted and removed, and chromatin bound to 6 µg of anti-SETDB1 antibody per condition at 4°C overnight followed by ChIP and elution in 30 µL of elution buffer using the Zymo ChIP DNA clean and concentrator (Zymo Research D5205). For qPCR, 1 uL of DNA from ChIP or input samples were used in a 10 uL reaction with the PowerUP SYBR Green Master Mix (Applied Biosystems A25742), for 40 cycles with a 30 sec annealing/extension at 60°C. Data were analyzed using GraphPad Prism with an unpaired two-tailed t-test.

### Nuclear imaging and analysis

Wild-type or mutant K562 cells were cultured overnight with coverslips coated with 1 mg/mL Concanavalin A. Cells were gently washed with 1x PBS, fixed with 3.7% formaldehyde for 20 minutes, permeabilized with 0.1% Triton X in 1x PBS for 5 minutes, blocked with 3% BSA/0.01% Triton X-100 in PBS for 30 min and stained with 1ug/uL Hoechst 33342 for 30 minutes. Images were captured using the Nikon Eclipse Ti2 inverted microscope (Nikon TI2-LA-FL-2) and analyzed using ImageJ (National Institutes of Health). Approximately 100 cells were counted per replicate per condition, for a total of 2 replicates. Data were analyzed using GraphPad Prism with an unpaired two-tailed t-test.

### RNA sequencing

We utilized Plasmidsaurus’ ultra-fast RNA sequencing and analysis services. 500,000 cells were spun, washed with PBS and resuspended in 100 uL Zymo DNA/RNA shield (Cat# R1100-50-E) for analysis by Plasmidsaurus.

### Declaration of generative AI and AI-assisted technologies in the writing process

During the preparation of this work the authors used ChatGPT to assist with writing of the manuscript, specifically its conceptual flow and clarity. After using this tool/service, the authors reviewed and edited the content as needed and take full responsibility for the content of the published article.

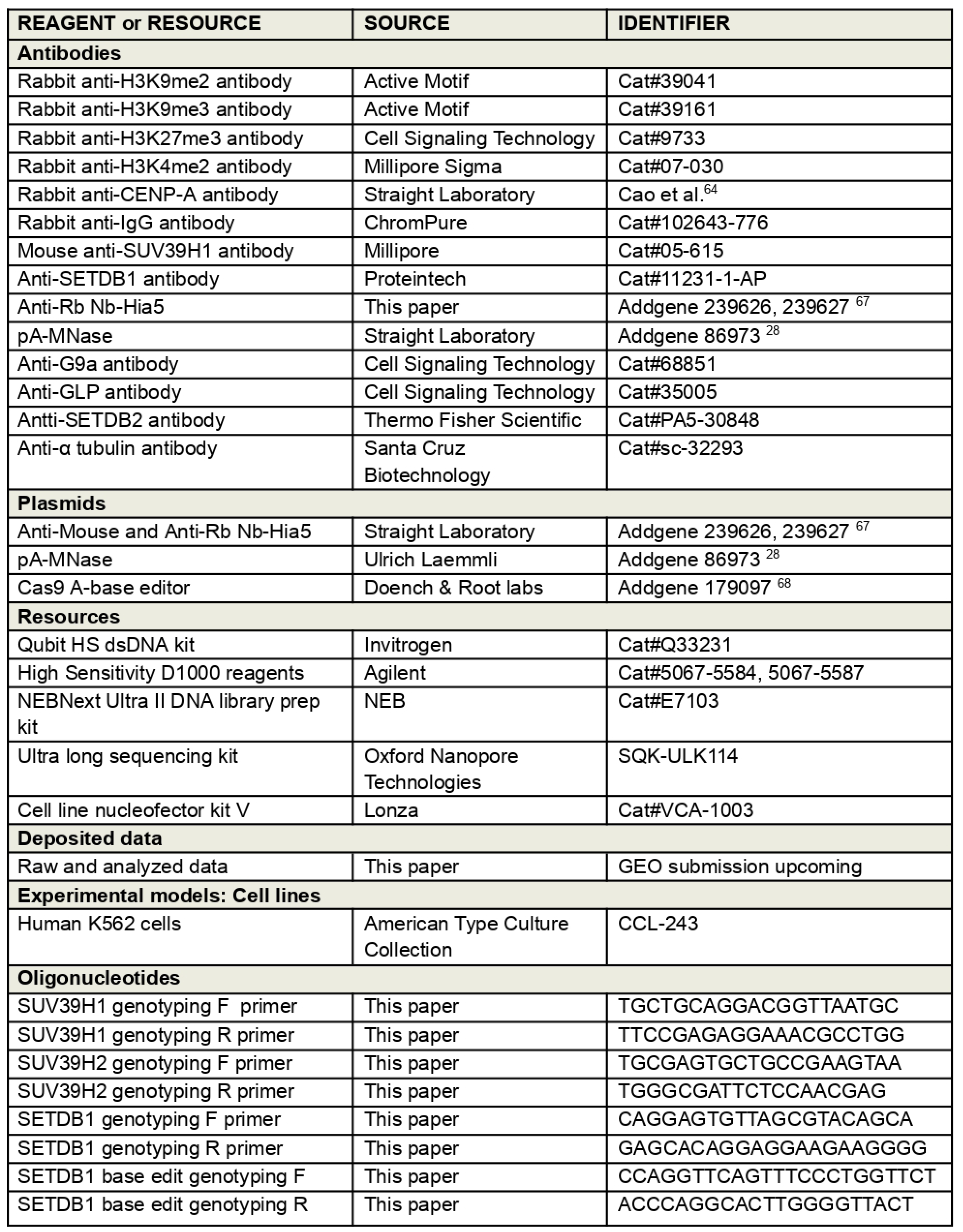
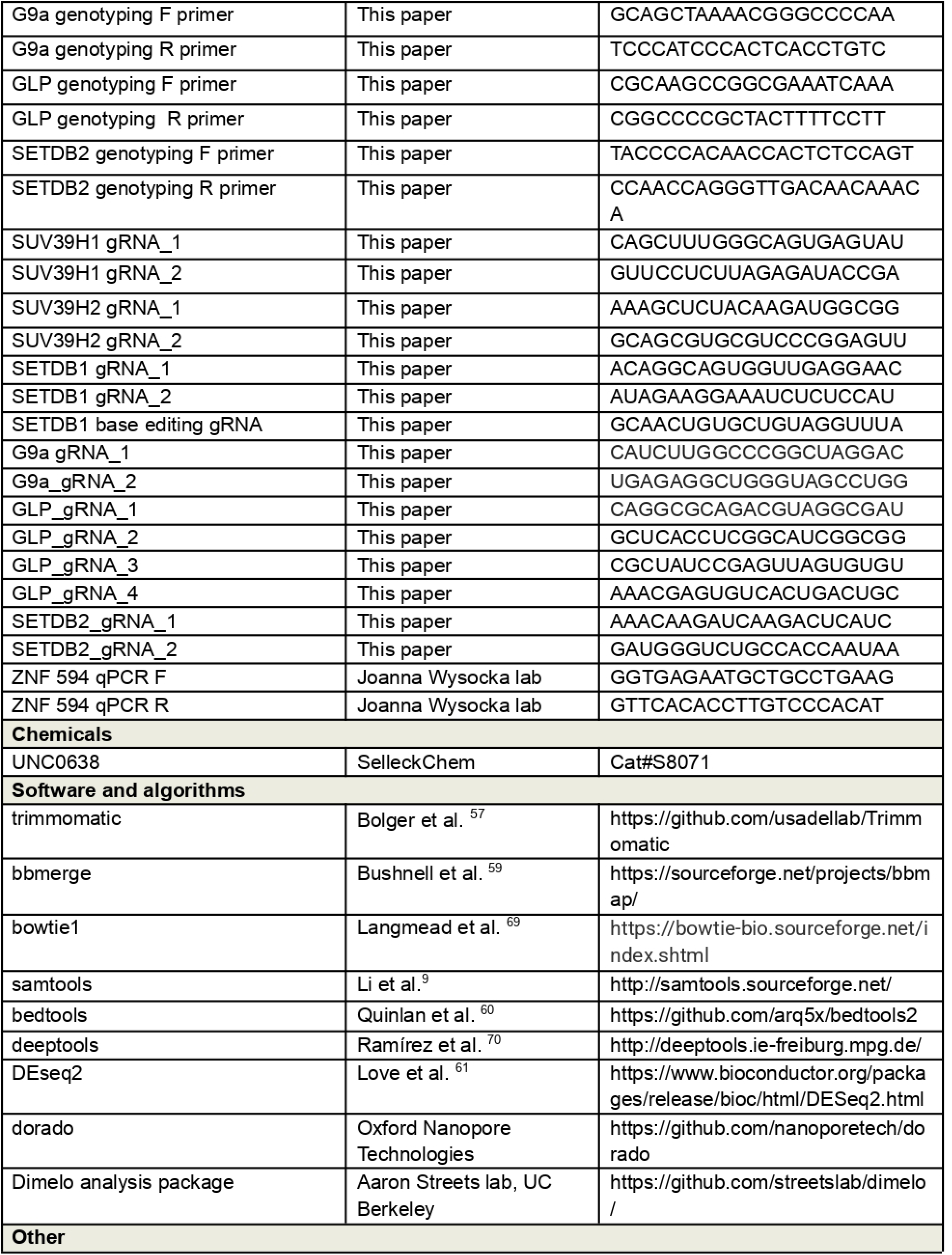
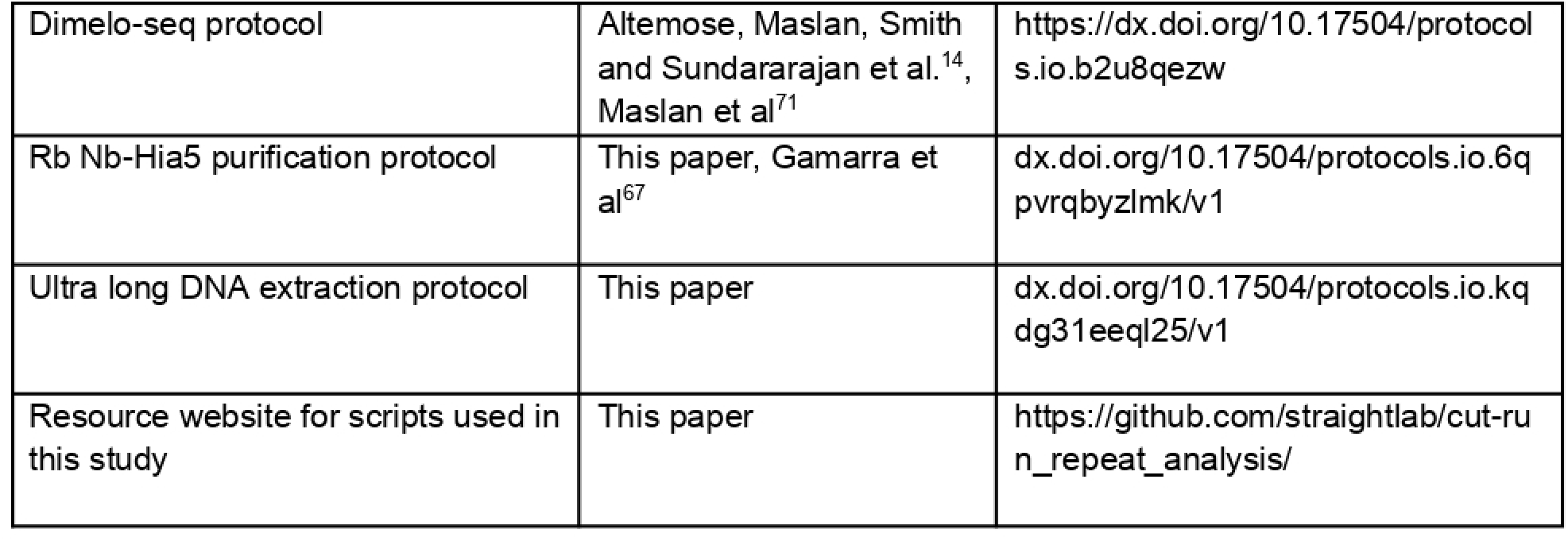

## Acknowledgements

The authors would like to thank Georgi K. Marinov for help with multimapping analysis and kindly providing access to python scripts and relevant resources, Marta Seczynska for helpful feedback on SETDB1 analysis, Shiran Bar for help with ChIP assays, and the members of the Straight lab for helpful discussion. This work was supported byNIH R01 GM074728 to AFS, Stanford School of Medicine’s Dean’s fellowship awarded to PS, AT is supported by NIH F32 GM165043, JPS and KAF were supported by NIGMS training grant T32 GM007276 and T32 GM007790 respectively.

---

**Figure S1:**
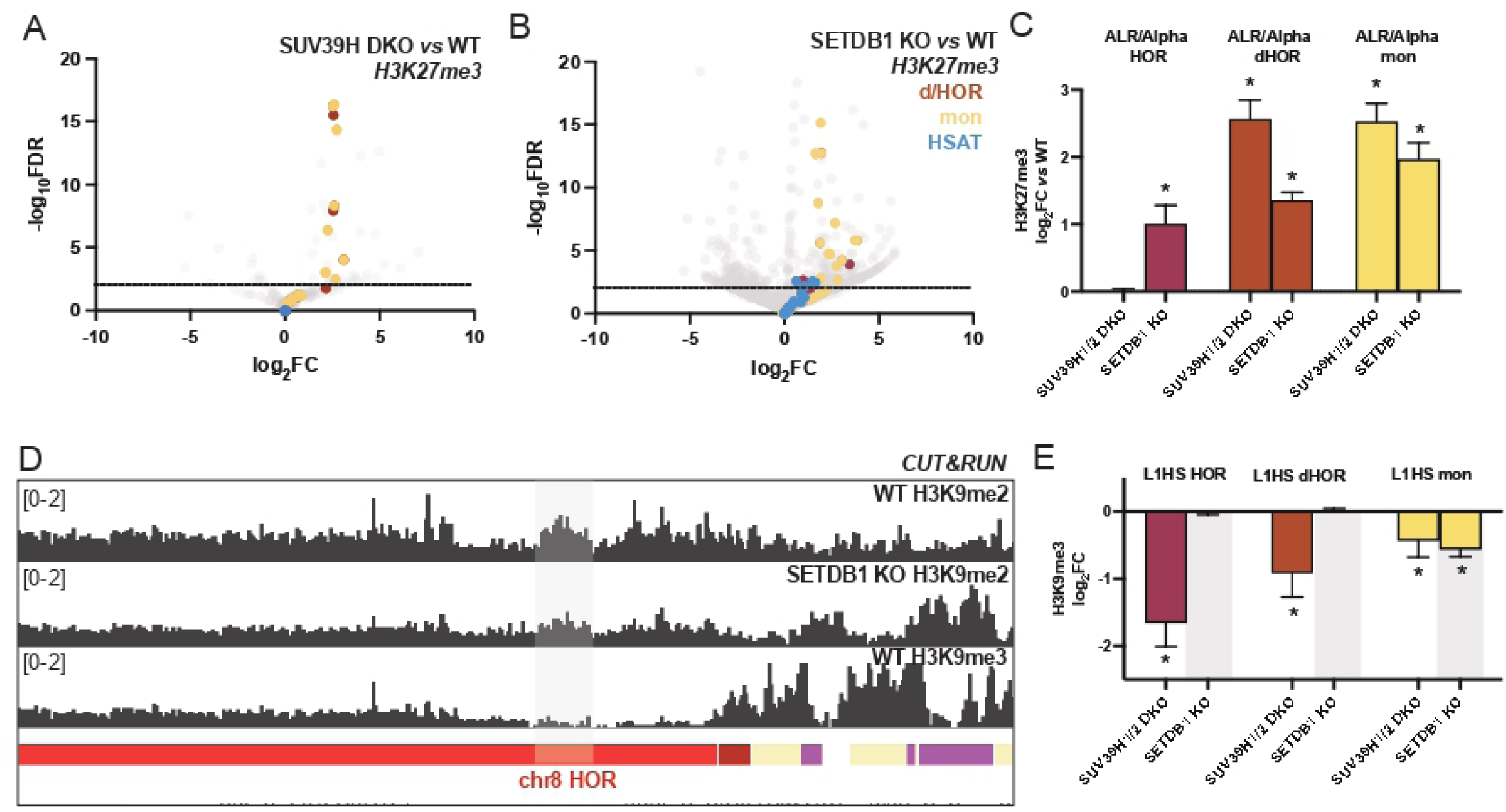
SUV39H and SETDB1 methyltransferases regulate facultative heterochromatin deposition. (A,B) Volcano plots of the the log_2_ fold change (log_2_FC) (X-axis) versus the -log_10_False Discovery Rate (-log_10_FDR) (Y-axis) for H3K27me3 at repeat elements located in HOR/dHOR (brown), monomeric repeats (yellow), and HSAT regions (blue), with 50,000 randomly sampled genomic or repetitive loci (grey). (A) SUV39H DKO compared to WT, (B) SETDB1 KO compared to wild-type. (C) Comparison of H3K27me3 log_2_FC versus WT in SUV39H DKO and SETDB1 KO at ALR/Alpha repeats embedded in HOR (magenta), dHOR (brown) and monomeric repeats (yellow). Error bars represent standard error of log_2_FC, asterisks represent padj<0.05. (D) IGV plot of the chromosome 8 HOR and adjacent region on the q-arm. The top two rows show relative levels of H3K9me2 in WT and SETDB1 KO and the third row shows comparison to H3K9me3 in WT. The gray bar highlights a SETDB1-dependent H3K9me2 peak that is absent in H3K9me3.(E) Comparison of H3K9me3 log_2_FC versus WT in SUV39H DKO and SETDB1 KO at L1Hs elements. Shown are ALR/Alpha repeats embedded in HOR (magenta), dHOR (brown) and monomeric repeats (yellow). Error bars represent standard error of log_2_FC, asterisks represent padj<0.05.

**Figure S2:**
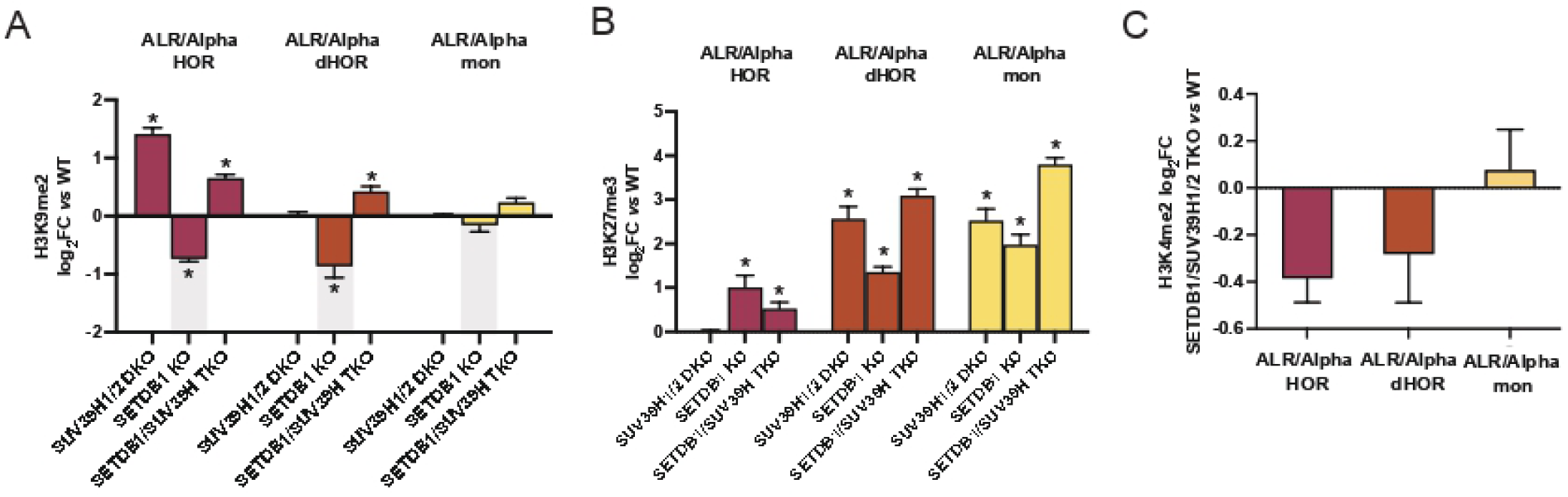
Epigenetic changes in SETDB1/SUV39H triple knockouts. H3K9me2 (A) and H3K27me3 (B) log_2_FC vs. wild-type for SUV39H1/2 DKO, SETDB1 KO and SETDB1/SUV39H TKOs at ALR/Alpha repeats embedded in HOR (magenta), dHOR (brown) and monomeric repeats (yellow). Error bars represent standard error of log_2_FC, asterisks represent padj<0.05. (C) H3K4me2 log_2_FC in SETDB1/SUV39H TKO vs. wild-type at ALR/Alpha repeats embedded in HOR (magenta), dHOR (brown) and monomeric repeats (yellow).

**Figure S3:**
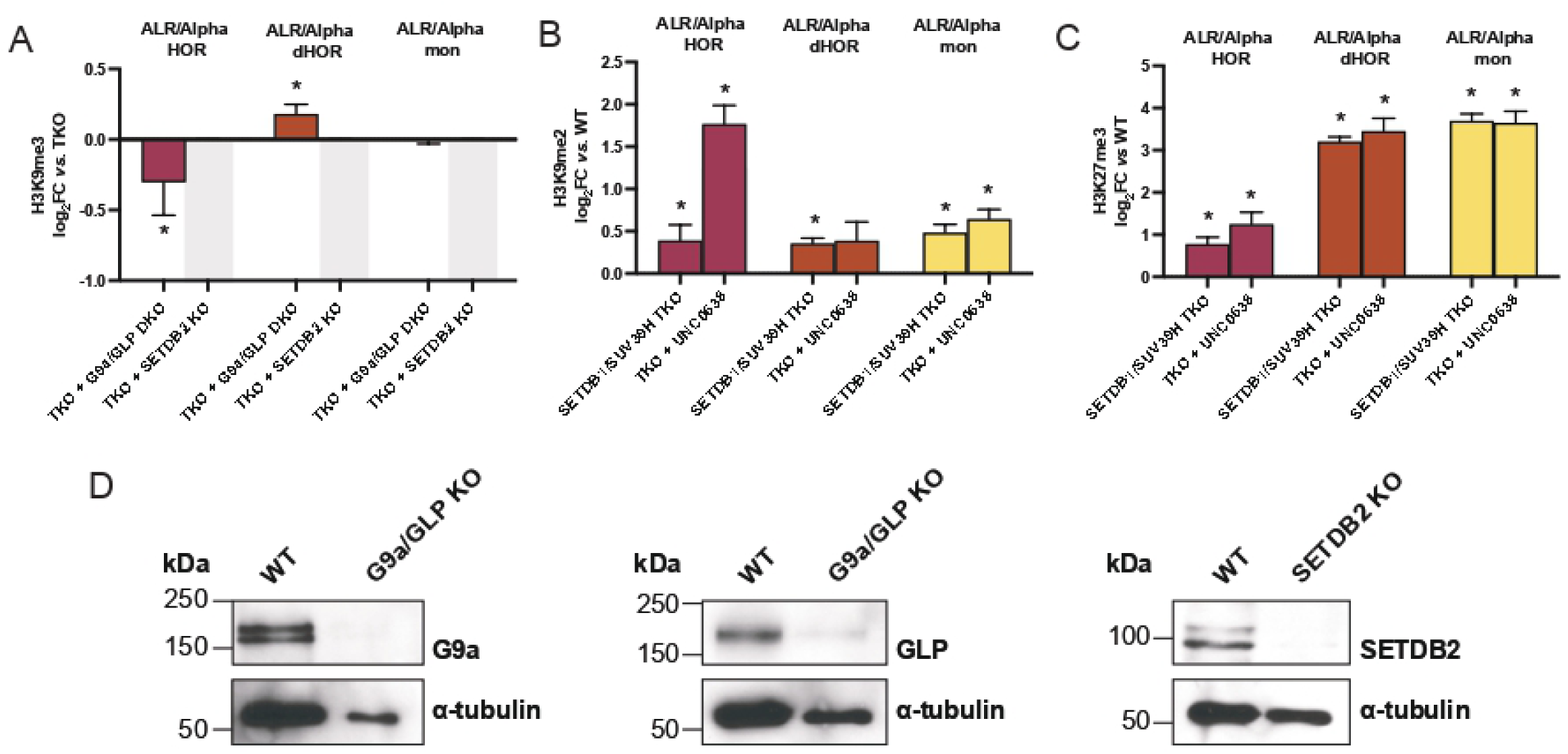
Heterochromatin regulation in SETDB1/SUV39H TKOs. (A) H3K9me3 log_2_FC in TKOs acutely depleted of G9a/GLP or SETDB2 compared to TKOs at ALR/Alpha repeats embedded in HOR (magenta), dHOR (brown) and monomeric repeats (yellow) Error bars represent standard error of log_2_FC, asterisks represent padj<0.05. (B-C) H3K9me2 and H3K27me3 log_2_FC vs. wild-type for SETDB1/SUV39H TKO at ALR/Alpha repeats in HORs (magenta), dHORs (brown) and monomeric repeats (yellow) show that TKOs have higher levels of these epigenetic marks at HORs as well. (D) Western blot analysis for G9a, GLP and SETDB2 following acute depletion of these methyltransferases as in (A) shows efficient knockout.

**Figure S4:**
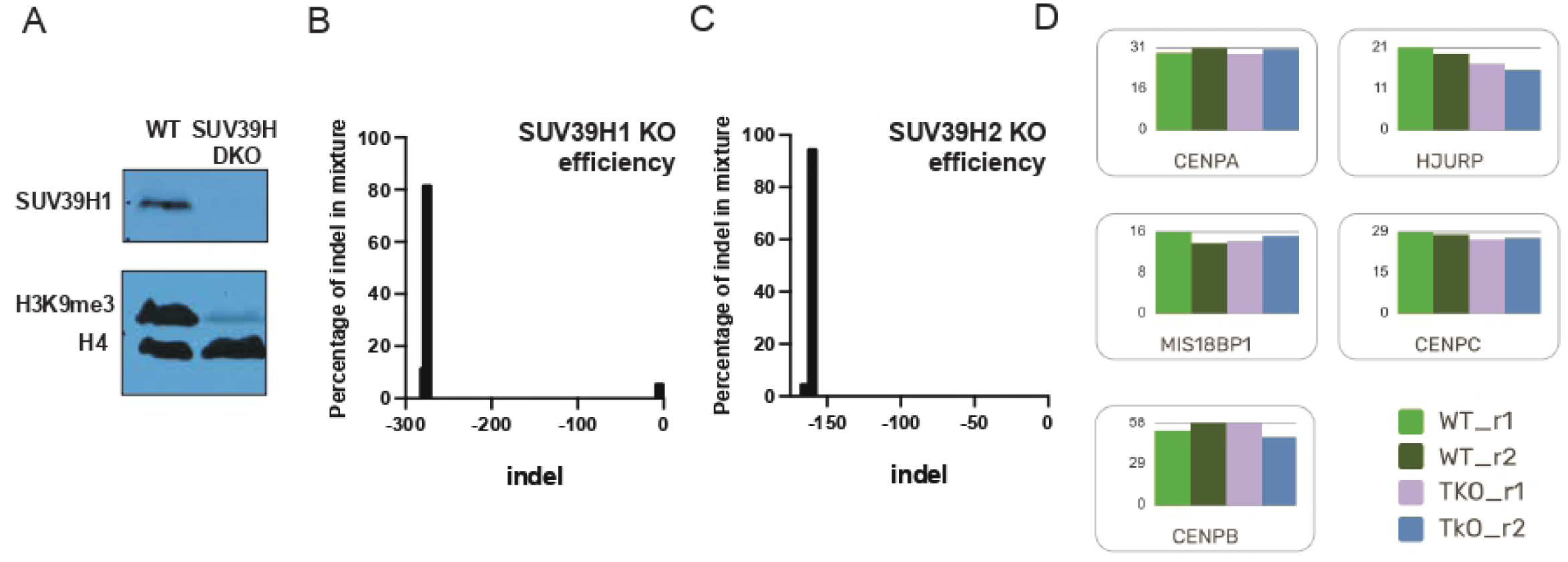
Characterization of SUV39H DKO and SETDB1/SUV39H TKOs. (A) Western blot analysis of SUV39H1 and H3K9me3 in wild-type and SUV39H DKO pools shows that both SUV39H1 and H3K9me3 are depleted in the SUV39H DKO. (B-C) Inference of CRISPR edits (ICE) analysis of the SUV39H1 locus (A) and SUV39H2 locus (B) shows high percentage of indels in SUV39H1 and SUV39H2 knockout pools. (D) RNA-sequencing analysis with counts-per-million on the Y-axis and condition on X axis shows that major centromere regulators are unchanged between WT and SETDB1/SUV39H TKOs.

**Figure S5:**
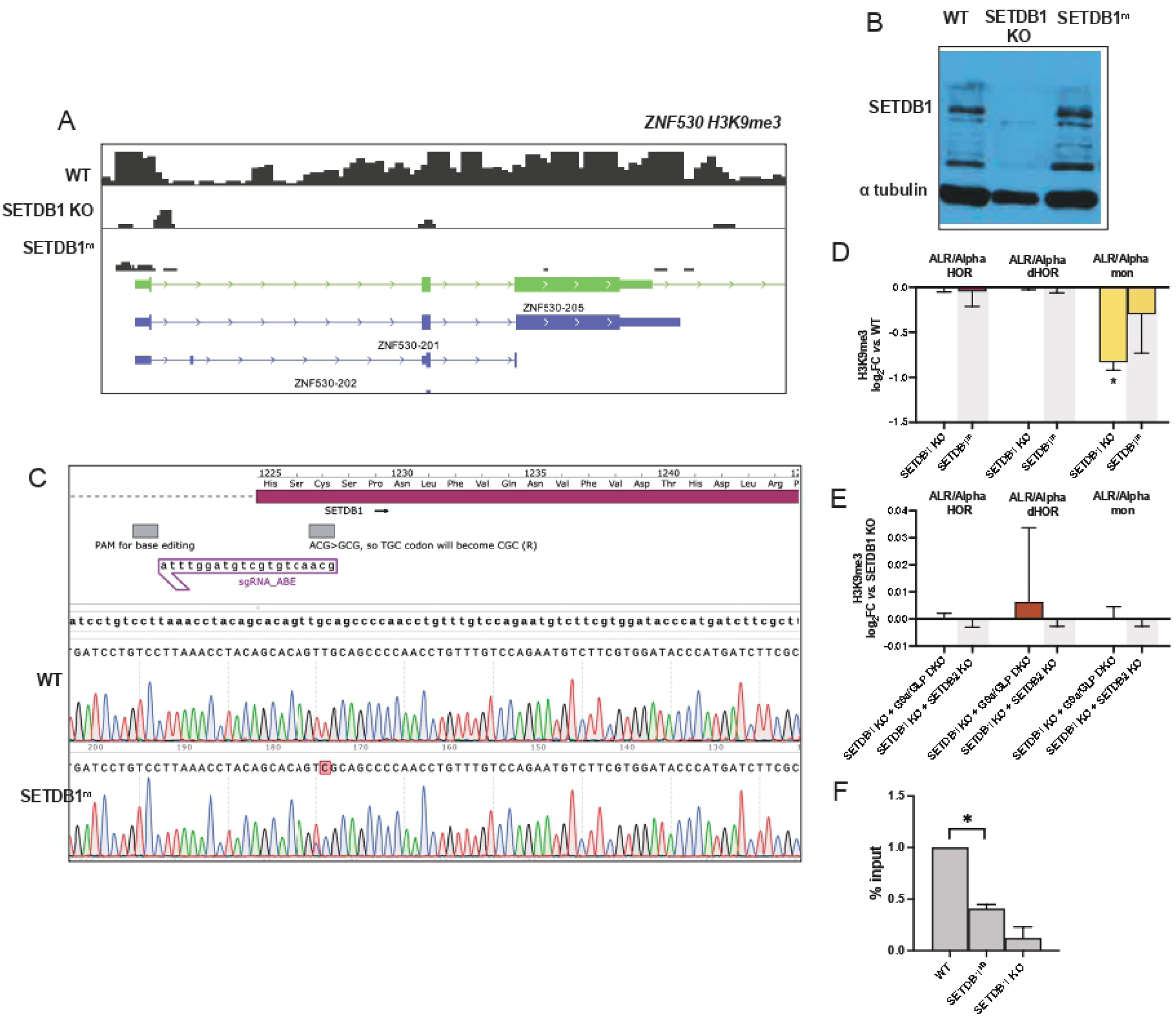
Characterization of SETDB1 KO and SETDB1^m^ catalytically inactive mutants. (A) H3K9me3 CUT&RUN coverage plot focusing on a ZNF530 locus on chr19 shows that both SETDB1 KO and SETDB1^m^ catalytically inactive mutant lose H3K9me3 at this locus. (B) Western blot analysis of SETDB1 in wild-type (WT), SETDB1 KO and SETDB1^m^ cells shows that SETDB1 is depleted in the KO but expressed at WT levels in SETDB1^m^ cells. (C) Sanger sequencing traces of the SETDB1 locus in WT (top) and SETDB1^m^ cells (bottom) shows a T>C change that translates into an arginine instead of a cysteine. Note that the T occupying the position next to it has also been edited in one of alleles; however this is the wobble position and does not change the corresponding amino acid. (D) H3K9me3 log_2_FC vs. wild-type for SETDB1 KO and SETDB1^m^ catalytically inactive mutant at ALR/Alpha repeats embedded in HOR (magenta), dHOR (brown) and monomeric repeats (yellow). Error bars represent standard error of log_2_FC. (E) H3K9me3 log_2_FC in SETDB1 KOs with acute depletion of G9a/GLP or SETDDB2 compared to. SETDB1 KO at ALR/Alpha repeats embedded in HOR (magenta), dHOR (brown) and monomeric repeats (yellow). Error bars represent standard error of log_2_FC. (F) ChIP-qPCR analysis using the percent-input method, normalized to WT levels (WT = 1), revealed significantly reduced SETDB1 occupancy at ZNF loci in SETDB1^m^ cells relative to WT controls (unpaired two-tailed t-test, p=0.0025).

**Figure S6:**
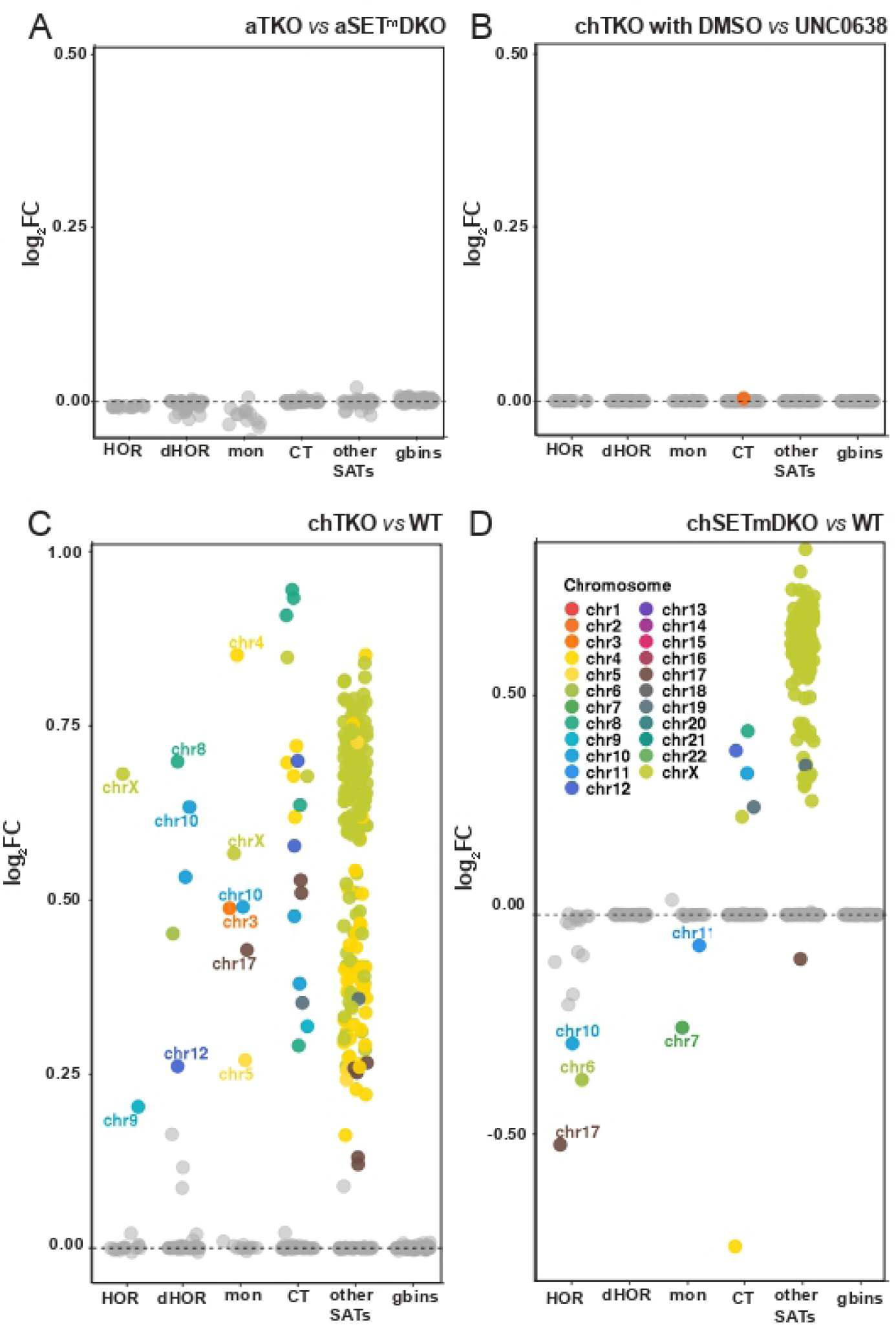
CENP-A spreading in multiple conditions. Dot plots showing CENP-A log_2_FC in acutely depleted SETDB1/SUV39H TKO pools relative to matched SETDB1^m^/SUV39H DKO (A), chronically depleted TKO pools treated with DMSO vs. UNC0638 (B), chronically depleted TKO pools compared to wild-type (C) and chronically depleted SETDB1^m^/SUV39H DKO pools compared to WT, across various repeat classes and genomic bins on individual chromosomes. Colored dots indicate padj < 0.05.

## Notes

https://github.com/straightlab/cut-run_repeat_analysis

### Competing Interest Statement

The authors have declared no competing interest.

### Summary of Updates

This revised version of the manuscript includes new data on the phenotypic effects of disruption of the SUV39H1/H2 and SETDB1 on chromosome segregation. The revision also includes updated data on the non-catalytic roles for SETDB1 in boundary maintenance and the relationship to DNA methylation. New chromosome by chromosome analysis of histone methylation and CENP-A levels are added to this manuscript. In addition, perturbation of G9A/GLP and SETDB2, new data on CENP-A spreading and RNA seq data characterizing the RNA changes in response to triple knockout of SUV39H1/H2 and SETDB1 have been included.

